# Polypyrimidine Tract Binding Protein blocks microRNA-­124 biogenesis to enforce its neuronal specific expression

**DOI:** 10.1101/297515

**Authors:** Kyu-Hyeon Yeom, Simon Mitchell, Anthony J. Linares, Sika Zheng, Chia-Ho Lin, Xiao-Jun Wang, Alexander Hoffmann, Douglas L. Black

## Abstract

MicroRNA-124 is expressed in neurons, where it represses genes inhibitory for neuronal differentiation, including the RNA binding protein PTBP1. PTBP1 maintains non-neuronal splicing patterns of mRNAs that switch to neuronal isoforms upon neuronal differentiation. We find that pri-miR-124-1 is expressed in mouse embryonic stem cells (mESCs) where mature miR-124 is absent. PTBP1 binds to this precursor RNA upstream of the miRNA stem-loop to inhibit mature miR-124 expression *in vivo*, and DROSHA cleavage of pri-miR-124-1 *in vitro*. This new function for PTBP1 in repressing miR-124 biogenesis adds an additional regulatory loop to the already intricate interplay between these two molecules. Applying mathematical modeling to examine the dynamics of this regulation, we find that the pool of pri-miR-124 whose maturation is blocked by PTBP1 creates a robust and self-reinforcing transition in gene expression as PTBP1 is depleted during early neuronal differentiation. While interlocking regulatory loops are often modeled between miRNAs and transcriptional regulators, our results indicate that miRNA targeting of posttranscriptional regulators also reinforces developmental decisions. Notably, induction of neuronal differentiation observed upon PTBP1 knockdown likely results from direct de-repression of miR-124, in addition to indirect effects previously described.

## Introduction

MicroRNAs (miRNAs) are an abundant class of molecules that regulate many important developmental events (Bushati and Cohen 2007). Assembled into an RNA induced silencing complex (RISC), miRNAs base pair with the 3’ untranslated regions (UTR) of their target messenger RNAs to inhibit translation and induce mRNA decay (Filipowicz et al. 2008; Braun et al. 2011; Chekulaeva et al. 2011; Tat et al. 2016). MiRNA biogenesis starts with transcription of a primary microRNA transcript (pri-miRNA) containing a hairpin structure that is cleaved by the DROSHA-DGCR8 (Microprocessor) complex to generate an ~ 70 nt stem-loop intermediate, the precursor miRNA (pre-miRNA). This pre-miRNA is exported to cytoplasm and further processed by the DICER-TRBP complex to produce a mature ~ 22 nt miRNA that is loaded onto an Argonaute protein within the RISC (Kim et al. 2009). Through base-pairing to its seed region, each miRNA mediates binding of RISC to miRNA responsive elements within a target mRNA to inhibit its expression. Besides the initial transcription of the pri-miRNA, miRNA expression can be regulated at later stages of biogenesis including the DROSHA and DICER processing steps (Rybak et al. 2008; Heo et al. 2009; Siomi and Siomi 2010).

The differentiation of embryonic cells into neurons is mediated by complex regulatory networks involving all the steps of gene expression pathway. Numerous molecules control this process, including transcription factors, chromatin modifiers, RNA binding proteins (RBPs) and miRNAs (Qureshi et al. 2010; Yoo et al. 2011; Hobert 2011; Kawahara et al. 2012; Sun et al. 2013; Staahl and Crabtree 2013; Vuong et al. 2016). The miRNA miR-124 has been described as a master regulator of neuronal differentiation (Visvanathan et al. 2007; Makeyev et al. 2007; Sun et al. 2013). MiR-124 is up-regulated as neuronal progenitors exit mitosis and begin to differentiate, and acts to repress many genes that maintain the non-neuronal state. Expression of miR-124 has been shown to be sufficient to drive cells into the neuronal pathway (Watanabe et al. 2004; Cheng et al. 2009; Yoo et al. 2011). Known targets of miR-124 include the RE1-silencing transcriptional factor (REST) that represses neuronal transcription programs, and the polypyrimidine tract binding protein (PTB, PTBP1 or hnRNP I) that represses neuronal alternative splicing patterns (Conaco et al. 2006; Visvanathan et al. 2007; Makeyev et al. 2007). In the human and mouse genomes, there are three genes encoding miR-124 precursors: pri-miR-124-1 (also called *Rncr3*), pri-miR-124-2 and pri-miR-124-3 (Blackshaw et al. 2004; Sanuki et al. 2011; Neo et al. 2014). In mouse, pri-miR-124-1 (chr14:64590657-64590741; mm9) and pri-miR-124-2 (chr3:17795662-17795770) are highly expressed in neurons, while expression of pri-miR-124-3 (chr2:180894040-180894107) is more limited. During neuronal development, the three loci show different levels and timing of induction (Deo et al. 2006; Cao et al. 2007; Sanuki et al. 2011).

PTBP1 represses neuronal patterns of alternative splicing in a variety of non-neuronal cell types including ESCs and neural progenitor cells (NPCs) (Shibayama et al. 2009; Keppetipola et al. 2012; Zheng et al. 2012; Xue et al. 2013; Linares et al. 2015). During neuronal differentiation, PTBP1 expression is turned off, in part through the action of miR-124 (Makeyev et al. 2007). This allows the induction of PTBP2, a paralogous protein which has different regulatory properties and allows the production of spliced isoforms specific to differentiating neurons (Boutz et al. 2007; Makeyev et al. 2007; Licatalosi et al. 2012; Zheng et al. 2012; Li et al. 2014; Gueroussov et al. 2015; Linares et al. 2015; Xue et al. 2016; Vuong et al. 2016). In addition to its role in splicing, PTBP1 is also found in the cytoplasm where it can antagonize the action of miR-124 by binding in the 3’ UTRs of transcripts such as *CoREST* and *SCP1* (Xue et al. 2013). Like the ectopic expression of miR-124, the depletion of PTBP1 can induce neuronal differentiation through effects on both splicing and translation (Xue et al. 2013). Although these two posttranscriptional regulatory programs, miR-124 induction and PTBP1 depletion, are each sufficient to drive differentiation, a systems-level understanding of how these regulatory circuits interact in controlling the commitment decision is lacking.

Here we show that PTBP1 directly represses miR-124 expression at the level of pri-miRNA processing. We find that pri-miR-124-1 RNA is expressed in mESCs without production of mature miRNA. We show that PTBP1 binds pri-miR-124-1 and blocks DROSHA cleavage in the nucleus. Through mathematical modeling, we find that this new regulatory loop connecting miR-124 levels with those of PTBP1, enforces a sharp regulatory transition during neuronal development as PTBP1 is depleted and miR-124 expression is increased.

## Results

### Primary miR-124-1 is expressed but not processed into mature miRNA in mouse embryonic stem cells

Examining the expression of pri-miR-124 by RT-PCR in mESC, N2a neuroblastoma cells and cultured mouse cortical neurons (mCtx), we detected strong expression of pri-miR-124-1 and pri-miR-124-2 (Figure 1A), but not of pri-miR-124-3 (data not shown) in cortical neurons. Notably, we observed expression of pri-miR-124-1 in mESCs (Figure 1A and Table 1). Similar expression was observed by RNAseq (see below). In contrast, mature miR-124 was abundant in neurons as expected, but only present at the limit of RT-qPCR detection in mESCs (Figure 1B and Table 1).

**Figure 1.**
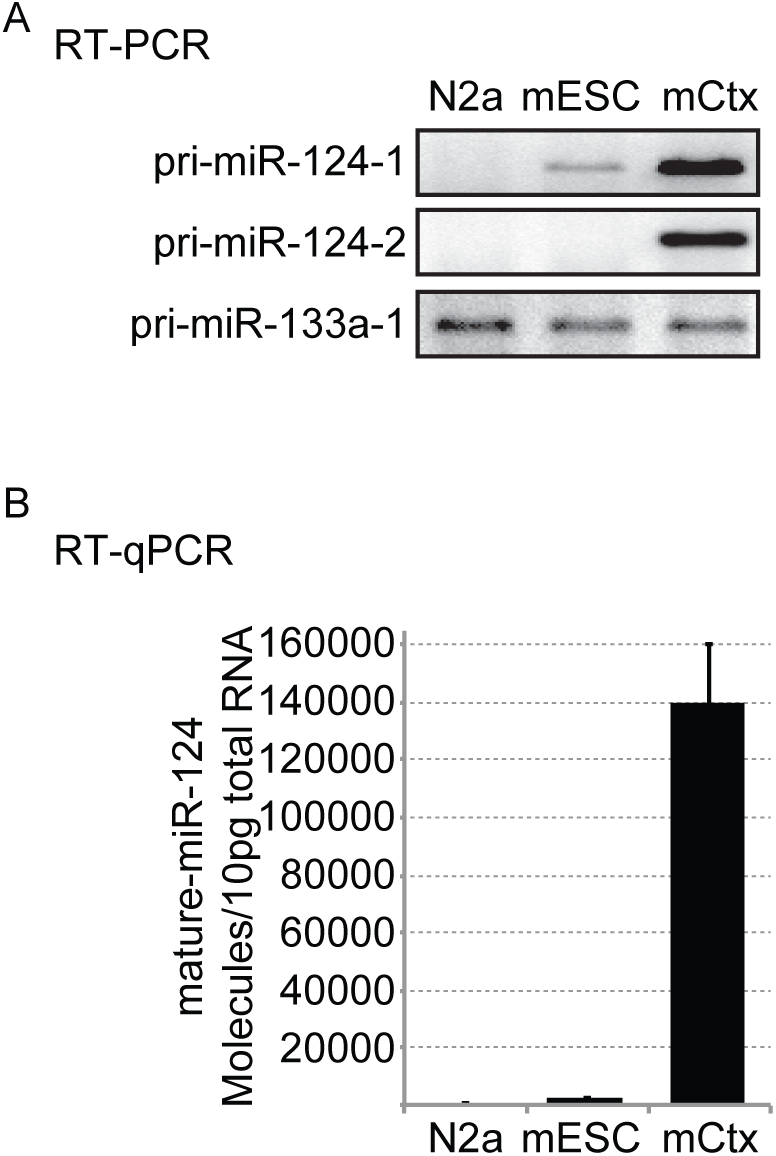
Pri-miR-124-1 but not mature miR-124 is expressed in mESCs. A. RT-PCR of pri-miR-124-1, pri-miR-124-2 and pri-miR-133a-1 from N2a, mESC and mCtx. Representative gel image is shown from 3 biological replicate cultures. RT-qPCR measurement of these molecules is reported in Table 1. N2a, mouse neuroblastoma cells; mESC, mouse embryonic stem cells; mCtx, mouse cortical neurons, day *in vitro* 5. B. Quantification of mature-miR-124 molecules in 10 pg of total RNA from the same cells as A. Mean molecule numbers were determined by RT-qPCR from 3 cultures; error bars are s.e.m.

**Table 1.** Quantification of primary and mature miR-124 and PTBP1/2. Molecules of RNA were determined by RT-qPCR and calculated estimating 10 pg of total RNA per cell. PTBP proteins were measured by western blot of protein from total cell lysate from a known number of cells and compared to a standard curve of recombinant His-PTBP protein. Total mass of protein per cell was measured to be ~ 0.4 ng (Supplemental Table 1) and used to estimate PTBP molecules per cell. Protein molecules from the ~ 0.4 ng of total protein were determined. N2a, mouse neuroblastoma cells; mESC, mouse embryonic stem cells; mNPC, mouse neuronal progenitor cells; mCtx, mouse cortical neurons. The mean ± s.d. of 3 cultures are given. Standard curves for RT-qPCR are presented in Supplemental Figure S1.

| Molecule |  | N2a | mESC | mNPC | mCtx |
| --- | --- | --- | --- | --- | --- |
| RNA | pri-miR-124-1 | 0.3 ± 0.1 | 3.6 ± 0.9 | 0.3 ± 0.2 | 30.1 ± 9.7 |
|  | pri-miR-124-2 | 0.0 ± 0.0 | 0.1 ± 0.0 | 9.7 ± 3.0 | 44.3 ± 11.7 |
|  | pri-miR-124-3 | 0.4 ± 0.3 | 0.8 ± 0.2 | 0.7 ± 0.1 | 3.1 ± 1.7 |
|  | mature-miR-124 | 3.1 ± 1.2 x10<sup>2</sup> | 2.2 ± 0.2 x10<sup>3</sup> | 1.3 ± 0.4 x10<sup>3</sup> | 1.4 ± 0.2 x10<sup>5</sup> |
|  | mature-miR-294 | 4.6 ± 4.5 | 2.4 ± 0.1 x10<sup>5</sup> | 11.6 ± 10.1 | 3.5 ± 6.0 |
|  | U6 | 2.5 ± 0.6 x10<sup>5</sup> | 2.1 ± 0.8 x10<sup>5</sup> | 2.5 ± 0.2 x10<sup>5</sup> | 2.7 ± 0.2 x10<sup>5</sup> |
| Protein | PTBP1 | 1.4 ± 0.4 x10<sup>4</sup> | 1.2 ± 0.2 x10<sup>4</sup> | 6.7 ± 2.2 x10<sup>3</sup> | 2.4 ± 8.8 x10<sup>2</sup> |
|  | PTBP2 | 4.6 ± 1.8 x10<sup>2</sup> | 2.3 ± 9.0 x10<sup>2</sup> | 8.5 ± 1.3 x10<sup>2</sup> | 3.5 ± 0.4 x10<sup>3</sup> |

Expression of miR-124 in mESCs was surprising given its role in driving neuronal differentiation. To better assess the amounts of mature miR-124 relative to its primary transcripts, we used RT-qPCR to quantify the absolute number of each RNA species in 10 pg of total RNA from each of the above cell types, as well as from isolated mouse neural progenitor cells (mNPCs). We determined that each of these cell types contain 10 to 19 pg of total RNA per cell (Supplemental Table S1), and that the different cell types expressed relatively constant levels of the U6 snRNA (Table 1). As expected, cortical neurons expressed between 60- and 500-fold more mature miR-124 than the other cell types. Similarly, mNPCs and N2a express about 100-fold lower levels of pri-miR-124-1 than neurons, although mNPCs have begun to express some pri-miR-124-2. Notably, the level of pri-miR-124-1 in mESCs was about 10-fold higher than in N2a cells and mNPCs, and only about 8-fold less than in cortical neurons (3.6 vs 30.1 molecules of pri-miR-124-1 per 10 pg of total RNA; Table 1). Comparing the level of mature miR-124 to the total of the three pri-miR-124 RNAs in the three cell types, we find that cortical neurons appear to convert about 4-fold more of the expressed pri-miR-124 into mature steady state miR-124, than do mESCs (Table 1; not accounting for miRNA turnover). Mature miR-294 was abundant in mESCs but not the other cell types, indicating that there was not a general loss of mature miRNAs in these cells. Repeated measurements in each cell type gave similar results indicating that the higher level of pri-miR-124-1 seen in mESCs was not due to random fluctuations or noise (Table 1). Thus, pri-miR-124-1 is expressed in mESCs, but the mature miRNA fails to accumulate due to either reduced processing or a higher decay rate.

### Pri-miR-124-1 is bound by PTBP1

Examining the sequences of the miR-124 genes, we noted a highly CU-rich segment upstream of the pri-miR-124-1 stem-loop that was not present in pri-miR-124-2 or pri-miR-124-3. This sequence is predicted to be bound by PTBP1, which we hypothesized may act to repress maturation of miR-124 in mESCs (Figure 2A) (Oberstrass et al. 2005; Han et al. 2014). Examining the miR-124-1 locus in an iCLIP dataset that we previously generated for PTBP1 in mESCs (Linares et al. 2015), we observed a significant peak of PTBP1 cross-linked fragments directly on this CU-rich segment of pri-miR-124-1, with a smaller peak of iCLIP tags upstream (Figure 2B). To confirm the interaction of PTBP1 with the pri-miRNA, we immunoprecipitated PTBP1 from mESC lysates, extracted the associated RNA, and performed RT-PCR for pri-miR-124-1 (Figure 2C). Indeed, pri-miR-124-1 was efficiently pulled down with PTBP1, and not in control immunoprecipitates (IPs). Pri-miRNAs not predicted to be bound by PTBP1 such as pri-miR-133a-1 and pri-miR-9-1 were not pulled down with PTBP1 IP. Other pri-miRNAs observed to be bound by PTBP1 in the iCLIP data such as miR-294/295 were also detected in our PTBP1 IPs but did not bind as efficiently as pri-miR-124-1 (Figure 2C and Supplemental Figure S2).

**Figure 2.**
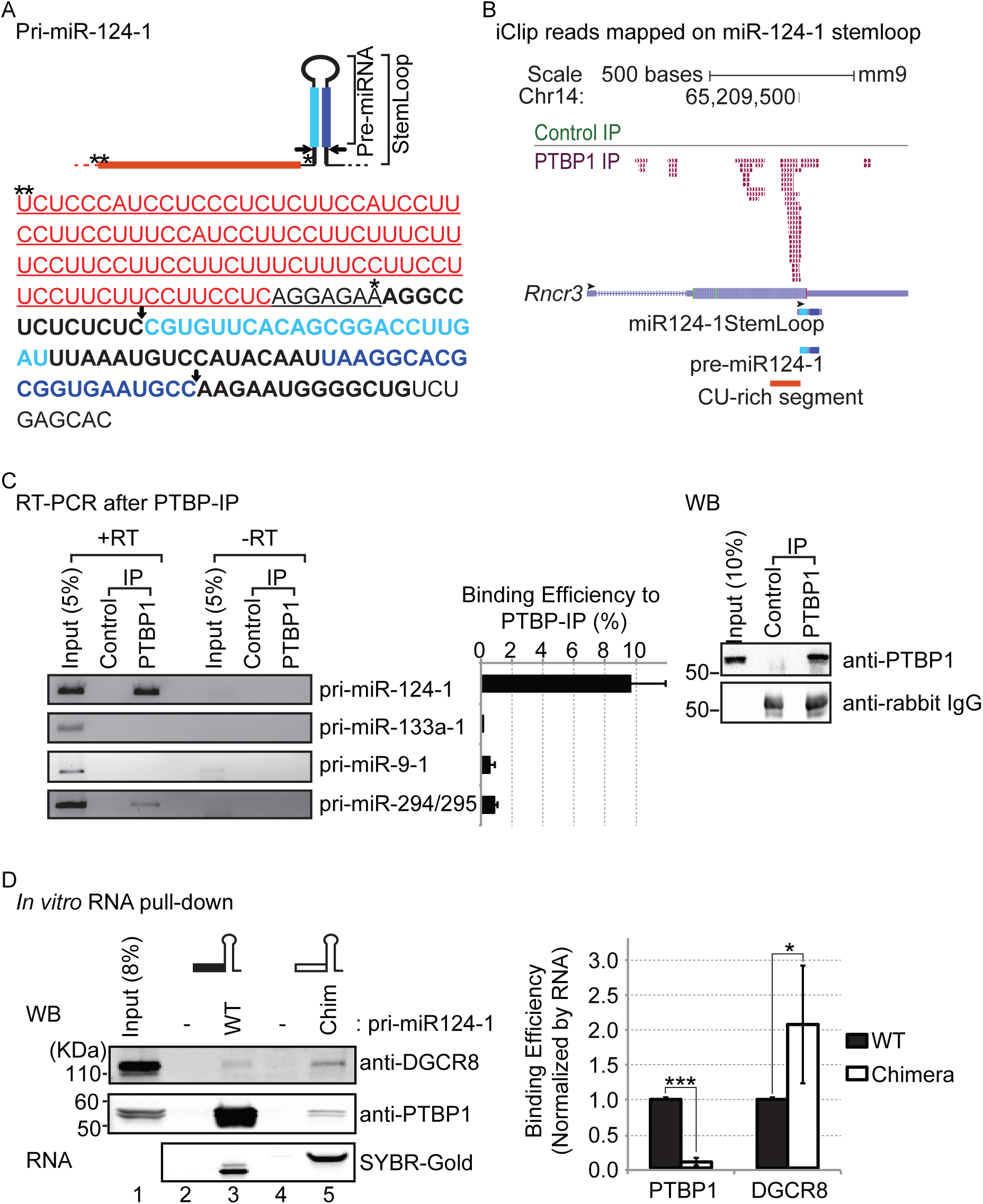
PTBP1 binds to a CU-rich segment in pri-miR-124-1. A. The sequence of pri-miR-124-1 with its secondary structure and sites of DROSHA cleavage (arrows) are diagrammed at top. The miR-124 duplex strands resulting from DROSHA and DICER cleavage are indicated in light and dark blue, with the final miR-124 in dark blue. Single and double asterisks mark the sequence locations of nucleotides 1 and 107 upstream from miR-124 stem-loop, respectively. The CU-rich segment extending from 107 nt to 8 nt upstream of the stem-loop of the miR-124-1 is in red. The sequences switched with pri-miR-1a-2 to form the chimeric RNAs in D are underlined. B. Genome browser view of PTBP1 iCLIP tags (PTBP IP, magenta) and FLAG-rabbit (Control IP, green) from mESCs aligning to the miR-124-1 host gene (*Rncr3, AK044422*). The stem-loop of miR-124-1, pre-miR-124-1 and the CU-rich segment are indicated at the bottom. Black arrowheads indicate 5’ to 3’ direction of each of RNA. C. Immunoprecipitation of pri-miR-124-1 with PTBP1. PTBP1 was isolated from mESC lysate by immunoprecipitation (PTBP1-IP) and bound RNA was extracted and assayed for pri-miRNAs by RT-PCR. An immunoprecipitate with rabbit IgG isotype was used as a negative control (Control-IP). RT-PCR products generated in the absence of reverse transcription (-RT) served as negative controls for genomic DNA contamination. Representative gel images from 3 biological replicates are shown at left. RNA binding efficiencies to PTBP1-IP were determined by comparison to 5 % of the input RNA (middle panel); error bars are s.e.m. Immunoprecipitated targets were checked by western blot (right panel). D. The CU-rich segment is required for the interaction between PTBP1 and pri-miR-124-1. A chimeric miRNA was generated by replacing the CU-rich segment of pri-miR-124-1 (black box) with the equivalent sequences of pri-miR-1a-2 (open box). Wildtype (WT) and chimeric (Chim) pri-miR-124-1 were transcribed *in vitro*, and hybridized to biotinylated adaptors immobilized on streptavidin beads. Beads carrying biotinylated adapters alone served as negative controls (lanes 2 and 4). Pri-miRNA bound beads were incubated with mESC total cell extract in 2 mM EDTA to inhibit DROSHA cleavage. After washing, bound PTBP1 and DGCR8 were detected by western blot and the bound RNA substrates were resolved on Urea-PAGE and stained with SYBR-Gold (left panel). Representative gel images from 3 biological replicates. Protein binding efficiencies to substrate RNAs were quantified and normalized to the amount of bait RNA and further normalized to WT RNA in each replicate (right panel). (n=3 biological replicates; Student’s t-test; * p≤0.05, *** p≤0.001; error bars are s.e.m.)

The CU-rich segment in pri-miR-124-1 extends from 107 nucleotides upstream of the miR-124-1 stem-loop to 8 nucleotides upstream (nt, - 107 to - 8). A small CU-rich segment is also present in the base-paired stem immediately 5’ of the DROSHA cleavage site (Figure 2A), and there are additional pyrimidine rich sequences approximately 200 nt upstream of the stem-loop. To examine the binding of PTBP1 to the CU-rich segment *in vitro*, we performed pull-down experiments using *in vitro* transcribed RNAs (Figure 2D). Wild type pri-miR-124-1 was compared to a chimeric RNA with nucleotides - 107 to - 1 upstream of the stem-loop replaced with the equivalent nucleotides from pri-miR-1a-2 (Figure 2A; underlined). These RNAs were hybridized to complementary biotinylated DNAs immobilized on streptavidin beads, incubated in mESC lysate, and the bound proteins then analyzed by western blot. As expected, PTBP1 bound efficiently to the wild type RNA and only minimally to the chimeric RNA, confirming its binding to the miR-124-1 upstream sequence (Figure 2D). Binding of the Microprocessor component DGCR8 exhibited a contrasting pattern, with greater binding to the chimeric substrate that is not bound by PTBP1 (Figure 2D).

To identify additional PTBP1 bound pri-miRNAs, we further analyzed the PTBP1 iCLIP data in mESCs (Supplementary Figure S3) (Linares et al. 2015). Defining sequence intervals from 125 nt upstream to 125 nt downstream of each annotated miRNA stem-loop, we identified 2020 PTBP1 iCLIP tags mapping within 295 pri-miRNA loci (Supplemental Figure S3A). In addition to those described above (Figure 2C), other pri-miRNAs showing significant PTBP1 binding included miR-5125, miR-7-1, miR-127 and others (Supplemental Figure S3). Some of these exhibited extensive PTBP1 binding outside of the defined search window (>125 nt upstream or downstream of the stem-loop). Such distal PTBP1 binding may not interfere with miRNA maturation, and it will be interesting to characterize the biogenesis of these miRNAs for possible PTBP1 regulation. Within sequences immediately adjacent to the miRNA stem-loops, pri-miR-124-1 had the highest number of PTBP1 iCLIP tags (53 total tags in the clusters described above; Figure 2B and Supplemental Figure S3A). Applying a predictive model of PTBP1 binding pri-miR-124-1 also yielded the highest PTBP1 binding score immediately upstream of the stem-loop of all miRNA loci (data not shown) (Han et al. 2014).

### PTBP1 inhibits miR-124-1 maturation

PTBP1 binding close to the site of pri-miR-124-1 processing indicated that the protein might interfere with DROSHA-DGCR8 binding and/or cleavage. To examine this, we compared the expression of mature miR-124 in *Ptbp1* wild type (WT) and knockout (KO) mESCs. A *Ptbp1* KO line was generated from a mESC line carrying loxp sites flanking *Ptbp1* exon 2, whose excision eliminates expression of the major PTBP1 isoforms PTBP1.1 and PTBP1.4. Cre recombinase was introduced into these cells and clones carrying homozygous deletions of the *Ptbp1* exon were selected (Figure 3A, lower panel). As observed previously, PTBP1 depletion induced the expression of PTBP2, which is encoded on a separate gene and is post-transcriptionally repressed by PTBP1 (Spellman et al. 2005; Boutz et al. 2007; Makeyev et al. 2007). A smaller, less abundant protein was also observed in the KO cells using an antibody targeting the C-terminal peptide common to both PTBP1 and PTBP2 (Markovtsov et al. 2000). This could be another isoform of PTBP2 or could result from translation initiation downstream from the deleted *Ptbp1* exon. Importantly, the loss of the major PTBP1 isoforms resulted in a nearly 5-fold increase in mature miR-124 over the control cells (Figure 3A; KO vs WT). Mir-294, whose precursor also exhibited some PTBP1 binding, showed only a small increase in the KO cells, while miR-9, which is not predicted to be targeted by PTBP1, showed no change.

**Figure 3.**
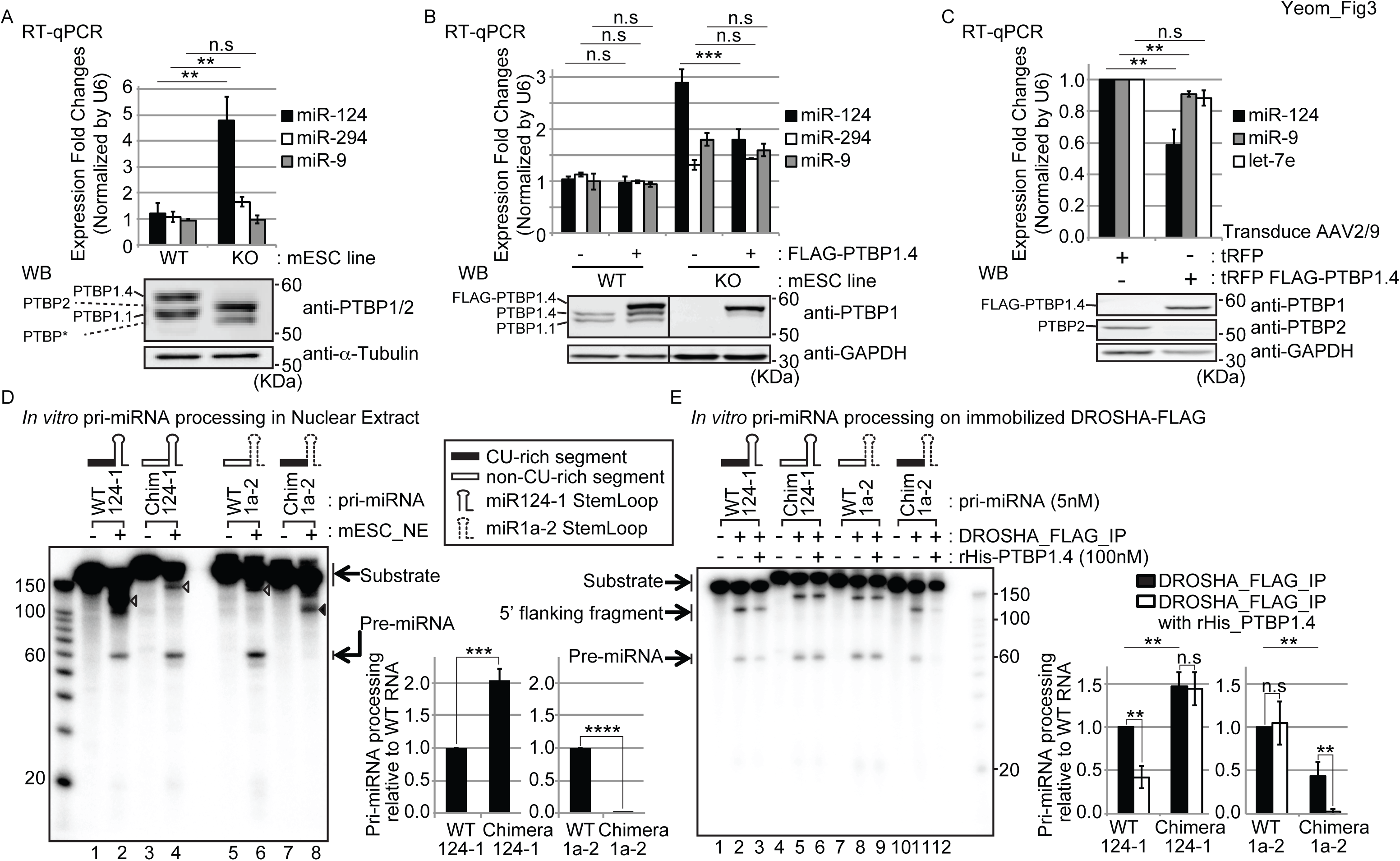
PTBP1 inhibits miR-124-1 maturation. A. Mature miR-124 is enhanced in a *Ptbp1* KO cell line. The levels of mature miR-124 were measured in wild type (WT) and *Ptbp1* knockout (KO) mESCs by RT-qPCR. miR-294 and miR-9 were assayed as control miRNAs (upper panel). All miR levels were normalized to U6 snRNA, and further normalized to the first replicate of WT cells. (n=3 cultures; Student’s t-test; ** p≤0.01; error bars are s.e.m.) *Ptbp1* depletion in the KO line was confirmed by western blot using a PTB_CT antibody targeting the common C-terminal peptide of PTBP1 and PTBP2 (lower panel) (Markovtsov et al. 2000). PTBP1 and PTBP2 isoforms are indicated to the left. Alpha-tubulin was used as a loading control. Representative gel images from 3 cultures. B. Ectopic PTBP1 reverses enhancement of miR-124 expression in the KO cell line. The level of mature miR-124 after ectopic FLAG-PTBP1.4 expression in WT and *Ptbp1* KO mESCs as measured by RT-qPCR. Mir-294 and miR-9 were assayed as control miRNAs (upper panel). All samples were normalized to U6 snRNA, and further normalized to the first replicate of WT cells with no transfected PTBP1. (n=3 biological replicates; Student’s t-test; *** p≤0.001; error bars, s.e.m.) Ectopic expression of PTBP1 (FLAG-PTBP1.4) was confirmed by western blot. GAPDH served a loading control (lower panel). Representative gel images from three replicates. C. Reintroduction of PTBP1 inhibits miR-124 expression in cultured cortical neurons. Cortical neurons were transduced with FLAG-PTBP1.4 expressing AAV one day after plating, and mature miRNA levels were measured at day 8 using RT-qPCR. Mir-9 and let-7e were assayed as controls (upper panel). Mature-miRNA level was normalized to U6 snRNA, and further normalized to the tRFP-only condition in each replicate. (n=3 biological replicates; Student’s t-test; ** p≤0.01; error bars are s.e.m.) Expression of FLAG-PTBP1.4 and endogenous PTBP2 were confirmed by western blot. GAPDH served a loading control (lower panel). Representative gel images from 3 replicates. D. The CU-rich segment inhibits pri-miRNA processing *in vitro*. Pri-miR-124-1, pri-miR-1a-2 and chimeric substrates (~ 3 nM) were incubated in mESC nuclear extract (NE) active for DROSHA processing and cleaved products were resolved by Urea-PAGE. For the chimeric RNAs the upstream sequences of miR-124-1 (Black box, containing the PTBP1 binding CU-rich segment) and miR-1a-2 (open box, non-CU-rich segment) were switched. The stem-loops of miR-124-1 and miR-1a-2 are indicated by solid and dashed lines, respectively. Un-processed substrates and the DROSHA processing products (pre-miRNAs) are indicated by arrows. Open arrowheads in lanes 2, 4 and 6 indicate the 5’ processing products (5’ flanking fragment) of pri-miR-124-1, chimera-miR-124-1 and pri-miR-1a-2 by DROSHA. The black arrowhead in lane 8 indicates an unidentified product of chimeric pri-miR-1a-2, whose size (~ 100nt) does not match any of the expected DROSHA cleavage products. Representative gel images from 3 replicates (left panel). Pri-miRNA processing efficiency was calculated by dividing pre-miRNA signal by un-processed substrate (incubated with buffer only), normalized by the number of labeled U residues in each substrate, and then to the WT construct in each replicate (right panel) (n=3 biological replicates; Student’s t-test; *** p≤0.001, **** p≤0.0001; error bars are s.e.m.) E. Recombinant His-PTBP1 inhibits processing of pri-miRNAs that have an upstream CU-rich segment. The same pri-miRNA substrates analyzed in D were subjected to *in vitro* pri-miRNA processing on immunoprecipitated DROSHA-FLAG with or without added rHis-PTBP1.4. Un-processed substrates and the DROSHA processing products (5’ flanking fragments and pre-miRNAs) are indicated by arrows. Representative gel images from 3 replicates (left panel). Pri-miRNA processing efficiency measured as in D is presented in the bar graph (right panel). (n=3 biological replicates; Student’s t-test; ** p≤0.01; error bars are s.e.m.)

**Figure 4.**
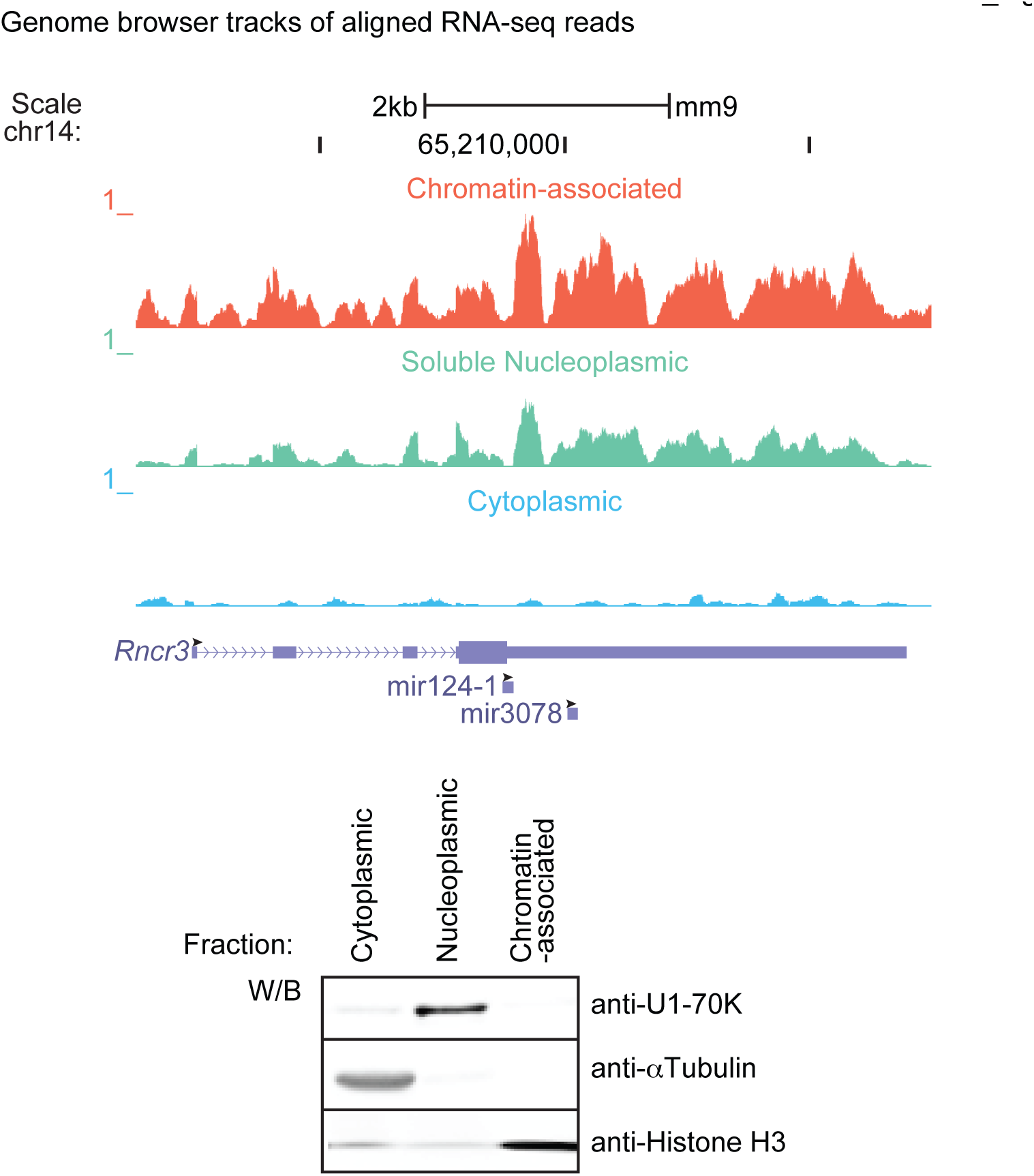
Pri-miR-124-1 is expressed in mESCs as chromatin-associated RNA but not as cytoplasmic RNA. Genome browser tracks showing RNA sequencing reads mapping to the pri-miR-124-1 host gene (*Rncr3, AK044422*) from three subcellular fractions: the chromatin-associated pellet (top), soluble nucleoplasm (middle) and cytoplasm (bottom). Fractionation followed the method of Wuarin and Schibler to enrich for nascent RNA in the chromatin fraction (Wuarin and Schibler 1994; Pandya-Jones and Black 2009; Yeom and Damianov 2017), and was assessed by western blot for separation of diagnostic proteins (lower panel; U1-70K for nucleoplasm, alpha-Tubulin for cytoplasm and Histone H3 for chromatin pellet). Read number scale (RPM) is set to 1 for all three fractions, the maximum peak height for the chromatin fraction. The positions of the two encoded miRNA stem-loops miR-124-1 and miR-3078 are indicated. Black arrowheads indicate the 5’ to 3’ direction of each of RNA. Tracks shown are the combined reads from 3 biological replicates (upper panel). Gel image is one of 3 biological replicates (lower panel).

We then introduced ectopic FLAG-PTBP1.4 into the WT and *Ptbp1* KO mESCs (Figure 3B). Ectopic PTBP1.4 had little effect on miR-124 or other miRNA levels in the WT cells where PTBP1 expression is already high. Under control transfection conditions, the KO showed a 3-fold increase in miR-124 expression over WT, with modest increases in miR-294 and miR-9. Reintroduction of PTBP1.4 reduced miR-124 in the KO cells by 40%, but had little effect on miR-294 or miR-9. From these data we conclude that PTBP1 inhibits expression of mature miR-124 in mESCs.

The induction of PTBP2 in the *Ptbp1* KO mESCs, and the coexpression of miR-124 and PTBP2 in neurons, indicate that PTBP2 does not repress miR-124 to the same degree as PTBP1. To examine whether miR-124 continues to be responsive to PTBP1 after neuronal differentiation, we transduced cultured cortical neurons with FLAG-PTBP1.4 using an adeno-associated virus (AAV). In these cells, miR-124 is expressed at high levels from both the miR-124-1 and miR-124-2 genes, while endogenous PTBP1 expression is repressed and replaced with PTBP2 (Table 1 and Figure 3C, lower panel). We found that re-expression of PTBP1 repressed miR-124 expression by 40%, corresponding to the proportion of miRNA arising from the pri-miR-124-1 gene (Table 1). The magnitude of the reduction in miR-124 also correlated with the titer of the transducing AAV (Supplemental Figure S4). Altogether, these data confirm that PTBP1 represses mature miR-124 expression *in vivo*.

The repression of miR-124 could occur by PTBP1 blocking DROSHA cleavage. However, most regulators of DROSHA processing have been found to bind in the stem-loop of the target RNA (Krol et al. 2010; Michlewski and Cáceres 2010; Ha and Kim 2014), while PTBP1 binds upstream of the stem. To test the effect of PTBP1 on DROSHA action, we used an *in vitro* pri-miRNA processing assay. We transcribed two pri-miRNA substrates: pri-miR-124-1 and pri-miR-1a-2, each including ~ 107 nt upstream and ~ 10 nt downstream of the miRNA stem-loop. We also designed two chimeric RNAs where the upstream regions of pri-miR-124-1 and pri-miR-1a-2 were switched. These RNAs were incubated in mESC nuclear extract active for DROSHA processing and their cleavage products characterized by gel electrophoresis. With incubation in nuclear extract, the synthetic pri-miR-124-1 was converted to the ~ 60 nt pre-miRNA with moderate efficiency, and a 130 nt species the size of the detached 5’ flanking fragment from upstream of the pre-miR-124-1 was also produced (Figure 3D, lane 2; open arrowhead). Using the chimeric RNA, where the PTBP1 binding site was replaced with the upstream sequence from miR-1a-2, production of the pre-miRNA increased two-fold (Figure 3D, lane 4). Note that the 5’ flanking fragment from the chimeric RNA yields a lower intensity band than the WT RNA due to its lower U content (Figure 3D). Interestingly, while the synthetic pri-miR-1a-2 was efficiently processed in the extract, this DROSHA cleavage was completely abolished by replacing its upstream segment with the PTBP1 binding sequence of miR-124-1 (Figure 3D, lanes 6 and 8; with bar graph to the right). Instead, we observed an aberrant product of ~100 nt that does not correspond in size to any of the expected DROSHA cleavage products (Figure 3D, lane 8; black arrowhead). These data indicate that the sequence upstream of the miR-124-1 stem-loop is inhibitory for cleavage at the expected DROSHA processing sites and may induce aberrant products from DROSHA or other activities.

To demonstrate that PTBP1 mediates the inhibition of DROSHA processing by the upstream CU-rich segment, we tested processing of the same four pri-miRNAs on immobilized DROSHA. We expressed Flag-tagged DROSHA in mES cells and pulled the protein down with anti-Flag antibodies. These IPs did not contain PTBP1 detectible by western blot (data not shown), and this system allowed clearer identification of the upstream processing products without other competing reactions. When incubated with the DROSHA-bound beads, all four pri-miRNAs were cleaved to produce the pre-miRNA, the 5’ flanking fragment and the 3’ flanking fragment (Figure 3E, lanes 2, 5, 8 and 11; The 3’ flanking fragments can be observed with longer exposure. Data not shown.). Processing of the two RNAs that lack a PTBP1 binding site (WT pri-miR-1a-2 and chimeric pri-miR-124-1) was not affected by added recombinant PTBP1.4 (Figure 3E, lanes 5,6 and 8,9; with bar graph to the right). In contrast, processing of both WT pri-miR-124-1 and the chimeric pri-miR-1a-2 containing the upstream PTBP1 binding site were strongly inhibited by the addition of PTBP1 (Figure 3E, lanes 2,3 and 11,12; with bar graph to the right). Taken together, these results demonstrate that PTBP1 inhibits DROSHA-DGCR8 processing of pri-miR-124-1 by binding to a CU-rich segment upstream of the stem-loop.

### The host gene for miR-124-1 is not expressed as an mRNA

The expression of pri-miR-124-1 in mESCs raises questions regarding the function of the transcript in these cells. The host gene for miR-124-1 (*Rncr3*) is annotated as a potential protein coding gene with an open reading frame that terminates within the miR-124 stem-loop, making the expression of the putative mRNA and the mature miRNA mutually exclusive. Thus, PTBP1 repression of DROSHA processing could allow expression of the host transcript as an mRNA. To examine this, we assessed the host gene transcripts that are spliced and exported to the cytoplasm in mESCs. As described previously, we isolated RNA from three subcellular fractions: nascent RNA that is associated with chromatin in the nucleus, RNA from the soluble nucleoplasm, and RNA from the cytoplasm where an mRNA will be translated (Pandya-Jones and Black 2009; Bhatt et al. 2012). We generated RNAseq datasets from each fraction and aligned the sequences to the genome to specifically examine reads mapping to the pri-miR-124-1 locus from these three compartments. A more complete analysis of these data will be presented elsewhere. A typical protein coding gene exhibits reads in the nuclear fractions derived from the exons and introns of the nascent RNA, and abundant exon reads with limited intronic reads in the cytoplasm derived from the spliced mRNA. This was not observed for the pri-miR-124-1 transcript, which expressed abundant reads for nascent RNA in the chromatin-associated fraction, similar to our observations by RT-qPCR (Table 1 and Figure 1A). RNA from the locus was also present at lower levels in the soluble nucleoplasm. However, reads from this gene were present at much lower levels in the cytoplasm and these reads were sporadically distributed without the typical exon peaks of a spliced mRNA. The lack of spliced RNA in the cytoplasm indicates that the host gene is not being expressed as an mRNA in mESC, although it could function as an mRNA in other cells.

### The regulation of miR-124 by PTBP1 enforces commitment to neuronal differentiation

We find that PTBP1 inhibits miR-124 expression in mESCs, while previous studies showed that miR-124 inhibits PTBP1 expression in neurons. One role for the early non-productive expression of miR-124 could be to alter the dynamics of the neuronal differentiation program. Since both miR-124 expression and PTBP1 depletion are known to induce neuronal differentiation, it was difficult to experimentally define a role for the PTBP1/miR-124-1 interaction through perturbation of the concentrations of each component. To examine how the antagonistic regulatory loops connecting PTBP1 and miR-124 affect their expression profiles over development, we constructed a kinetic computational model of miR-124 and PTBP1 production (Figure 5A). The model incorporated the known regulatory loops affecting PTBP and miR-124 expression, and was assessed with or without the addition of the newly discovered PTBP1 inhibition of miR-124-1 processing (Figure 5A; red line).

**Figure 5.**
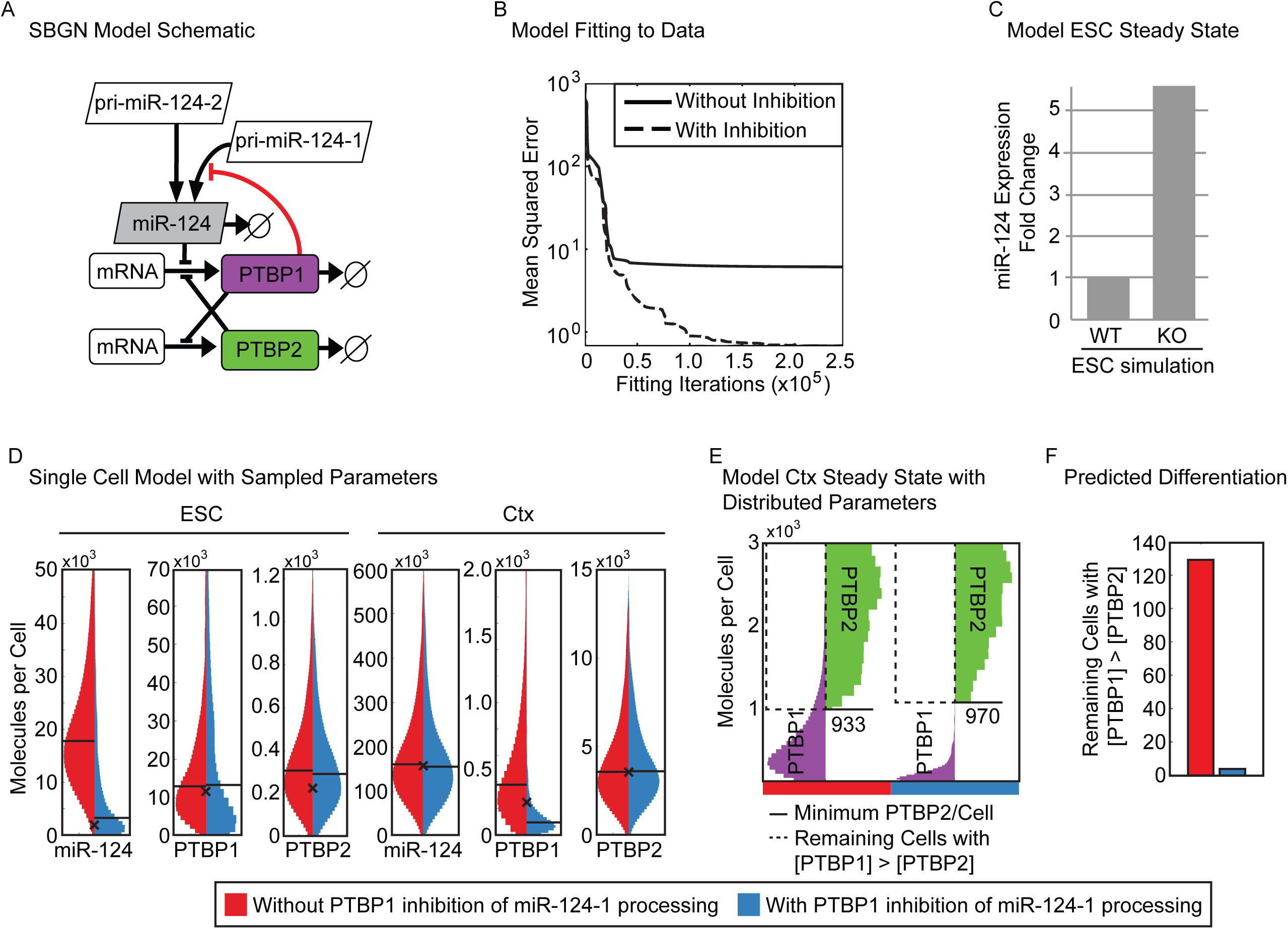
PTBP1 inhibition of miR-124-1 maturation mitigates cell-to-cell variability during neuronal differentiation. A. Schematic of the ordinary differential equation model of miR-124 expression presented in Systems Biology Graphical Notation (SBGN (Le Novère et al. 2009)). Known regulatory interactions are indicated with black lines and the inhibition of miR-124 processing by PTBP1 identified here is indicated by a red line. B. Mean squared error between best simulation fits and the molecule numbers measured experimentally (Table 1). Parameter estimation was performed by particle swarm optimization in either the model with the inhibition of pri-miR-124-1 processing by PTBP1 (dashed line) or without the addition of this mechanism (solid line). C. Prediction of miR-124 levels in the WT and PTBP1 KO ESC from simulations maintaining or removing the synthesis of PTBP1. D. Distribution of mature miR-124, PTBP1 and PTBP2 levels in embryonic stem cells (ESC) and cortical neurons (Ctx) in 1000 single-cell simulations run with (right, blue) and without (left, red) PTBP1 inhibition of pri-miR-124-1 processing. The experimentally measured value is indicated with an x and the median molecule numbers are indicated with horizontal lines. E. PTBP1 and PTBP2 molecules in 1000 simulated single cells of the Ctx stage at the end of the modeled time-course, with (right side, blue bar) and without (left side, red bar) PTBP1 inhibition of pri-miR-124-1 processing (shown in Supplemental Figure S5). The minimum PTBP2 molecule number is indicated with a solid line and those cells with more PTBP1 than this number are indicated in the region marked with a dashed line. F. Number of cells from 1000 single cell simulations in which the concentration of PTBP1 in the neuronal population was above the minimum PTBP2 concentration at this stage with (blue) and without (red) PTBP1 inhibition of pri-miR-124-1 processing (Number of cells within the region marked with a dashed line in E).

To provide input parameters for the model, we used the above-measured per-cell levels of each RNA species in each of three cell types: mESC, mNPC prior to the induction of miR-124, and cultured cortical neurons (mCtx) (Table 1). We also performed quantitative fluorescent western blots of PTBP1 and PTBP2, compared to a standard curve of recombinant protein, to measure protein molecules per cell (Table 1 and Supplemental Table S1). These concentrations were used to parameterize a system of ordinary differential equations describing the concentrations of all proteins and miRNAs determined by their rates of synthesis and decay, and the inhibition of these processes. The entire space of kinetic constants affecting miR-124 and PTBP1 concentrations was then scanned over 12 orders of magnitude to capture all potential parameter combinations (Figure 5B). Parameters were iteratively fit to the models to identify combinations that minimized the distance to experimentally observed concentrations.

We found that incorporating the inhibition of pri-miR-124-1 processing by PTBP1 into the model enabled a substantially better fit to the experimental data than was possible with a model lacking this inhibition (Figure 5B; dashed line). In fact, the model without PTBP1 repression of pri-miR-124-1 was unable to recapitulate the experimentally determined levels of protein and RNA with any parameterization, suggesting the experimental data cannot be quantitatively explained without this mechanism (Figure 5B; solid line). To assess how well the model with PTBP1 repression of miR-124-1 recapitulated the underlying conditions, we removed expression of PTBP1 to predict the effect of knocking out PTBP1 (a condition not used to parameterize the model). This simulation produced an approximately 5-fold increase in mature miR-124 upon loss of PTBP1 – notably close to the ~ 5-fold increase we observed in *Ptbp1* KO mESCs (Figures 5C and 3A). We conclude that the inhibition of miR-124-1 processing by PTBP1 is required to explain the quantitative experimental data and plays a substantial role in determining the balance of these molecules *in vivo*.

Feedback loops involving miRNAs and transcription factors add robustness to differentiation pathways (Tsang et al. 2007; Ebert and Sharp 2012; Vera et al. 2013; Posadas and Carthew 2014). By reducing the impact of variation in environment and genotype, these regulatory circuits make developmental transitions more reliable. We wanted to examine whether similar properties would result from regulatory loops involving miRNAs and RBPs. To test the effects of miR-124-1 regulation by PTBP1 on the path of neuronal differentiation under conditions of heterogeneous environments or intrinsic variability, we added stochastic parameter sampling to the kinetic models. Parameter sampling has been widely used to model cellular heterogeneity (Shokhirev et al. 2015; Yao et al. 2016; Mitchell and Hoffmann 2018). Simulations of 1000 individual cells were run for each model with each parameter sampled from a 4-fold range distribution centered on the optimal parameter identified by the original fitting to experimental data (Figure 5D). We found that the addition of feedback not only reduced the overall level of mature miR-124 in ESCs, but it also reduced cell to cell heterogeneity in this level (Figure 5D, left-most panel; miR-124 level in ESC, Red vs Blue distribution). Interestingly, we found that heterogeneity in PTBP1 levels was also greatly reduced in cortical neurons when miR-124-1 was repressed by PTBP1 (Figure 5D, fifth panel; PTBP1 levels in Ctx, Red vs Blue distribution). The tighter regulation of PTBP1 in the model affects the transition to PTBP2 expression in the presence of biological variability, with fewer cells reaching a state of incompletely down-regulated PTBP1, and ensuring that a neuronal, PTBP2-only state is reached (Figures 5E and 5F). Thus, the computational modeling indicates that PTBP1-meditated inhibition of miR-124-1 can act to mitigate biological variability and ensure that robust differentiation occurs in all cells.

## Discussion

### A posttranscriptional feedback loop affecting neuronal differentiation

Neuronal differentiation from progenitor cells involves a complex regulatory program affecting all levels of the gene expression pathway. Notch signaling through its receptor Jagged maintains the neuronal progenitor cell population and must be turned off to initiate differentiation (Louvi and Artavanis-Tsakonas 2006). The transcriptional repressor REST and the c-terminal phosphatase SCP1 are down-regulated to allow induction of neuron-specific genes (Ballas et al. 2005; Yeo et al. 2005), and the transcription factors Sox9 and histone methyltransferase Ezh2 that regulate glial cell fate are also repressed (Neo et al. 2014). As neuronal progenitors exit mitosis and begin to form early neurons, chromatin remodeling complexes acquire new subunits, including BAF53b in place of BAF53a to form the nBAF complex (Yoo et al. 2009). At the same time PTBP1 is down-regulated, inducing expression of the neuronal paralog PTBP2 and shifting the cells to a neuronal program of alternative splicing (Boutz et al. 2007; Makeyev et al. 2007; Licatalosi et al. 2012; Li et al. 2014). In addition to these general regulators, there are a large number of localized signaling pathways and specialized transcription factors that drive development of specific neuronal subtypes and lineages (Kessaris et al. 2014; Guo and Anton 2014; Bandler et al. 2017). As a master regulator of neuronal cell fate, miR-124 is induced at the beginning of NPC differentiation into neurons and directly represses expression from the *Jagged*, *REST*, *SCP1*, *BAF53a*, *Sox9*, *PTBP1* and many other mRNAs.

Of the three MiR-124 loci, only pri-miR-124-1 has been characterized genetically and was found to be essential for brain and retinal development (Blackshaw et al. 2004; Sanuki et al. 2011). MiR-124 expression is modulated by feedback loops where particular target molecules can alter miR-124 function or expression. For example, all three loci contain REST binding sites, which may limit their transcription in non-neuronal cells (Conaco et al. 2006). In other studies, 3’UTR binding by PTBP1 was found to block miR-124 action on its targets *RCOR1* (CoREST) and *SCP1* (Xue et al. 2013). These additional regulatory loops can repress miR-124 activity prior to neuronal differentiation.

We identify a new feature of the complex regulatory circuit controlling miR-124 that acts on its biogenesis rather than its transcription or functional activity. PTBP1 binding to pri-miR-124-1 blocks DROSHA cleavage and prevents formation of the pre-miRNA. PTBP1 has long been known to be repressed by miR-124 (Makeyev et al. 2007), and knockdown of PTBP1 is sufficient to induce neuronal differentiation in non-neuronal cell lines (Xue et al. 2013). This action was attributed to the loss of PTBP1 allowing miR-124 to then target *CoREST* and *SCP1* (Xue et al. 2013). However, our results indicate that an additional effect of PTBP1 depletion is the direct up-regulation of mature miR-124 itself. This may be the primary driver of the neuronal differentiation seen with PTBP1 knockdown.

The block to miR-124 biogenesis by PTBP1 reinforces the antagonistic regulatory interactions of these two molecules. Feedback loops between transcription factors that drive development of particular cell lineages and miRNAs that regulate mRNAs in that lineage have been described (Ebert and Sharp 2012; Posadas and Carthew 2014). In these systems, it is proposed that the feedback improves efficiency of commitment to differentiation and reduces noise. We find that the new PTBP1/miR-124-1 feedback loop alters the dynamics of neuronal differentiation, enforcing PTBP1 expression prior to differentiation and miR-124 afterwards, and making the cell fate decision more reliable in the presence of biological variability. This is similar to what is proposed for transcriptional feedback loops but here involves only posttranscriptional steps of gene expression (Tsang et al. 2007; Herranz and Cohen 2010; Ebert and Sharp 2012).

### The regulation of miRNA biogenesis

There are multiple examples of RBPs regulating miRNA processing, and recent studies identify widespread interactions between RBPs and stem-loop RNAs (Treiber et al. 2017; Nussbacher and Yeo 2018). In early development, Lin28 binds to the terminal loop of the let-7 miRNA, blocking DROSHA processing in the nucleus (Newman et al. 2008; Viswanathan et al. 2008) and DICER processing in the cytoplasm (Rybak et al. 2008; Heo et al. 2008; 2009). In other cells, the same let-7 stem-loop is bound by the KSRP and hnRNPA1 proteins that enhance or repress let-7 processing, respectively. HnRNPA1 also binds the terminal stem-loop of miR-18a, where it stimulates DROSHA cleavage (Guil and Cáceres 2007; Michlewski et al. 2008; Trabucchi et al. 2009; Michlewski and Cáceres 2010). The biogenesis of miR-7 is also regulated at the level of DROSHA processing. MiR-7 is primarily expressed in brain and pancreas, but the miR-7-1 stem-loop is within an intron of the widely expressed hnRNPK transcript (Landgraf et al. 2007; Hsu et al. 2007). Proteins implicated in repressing pri-miR-7-1 processing in non-neuronal cells include HuR and Musashi-2, again by binding to the terminal loop of pri-miR-7-1 (Choudhury et al. 2013). The regulation of miR-124-1 by PTBP1 differs from the above systems in that an extended inhibitory binding site is upstream from the stem-loop structure. A recent study identified the Quaking 5 protein (QKI5) as a positive regulator of pri-miR-124-1 processing during human erythropoiesis (Wang et al. 2017). The QKI5 binding site is upstream of the PTBP1 sites identified here and the role of PTBP1 in erythropoiesis has not been explored.

The CU-rich PTBP1 binding segment that inhibits DROSHA-DGCR8 processing extends nearly to the base of the stem where DROSHA cleavage occurs (Figure 2A). Thus PTBP1 may block access of DROSHA-DGCR8 to its cleavage site. Alternatively, PTBP1 binding may change the folded structure of pri-miR-124-1 to disrupt DROSHA-DGCR8 recognition. Interestingly, we observe reduced DGCR8 binding to the pri-miRNA in the presence of the upstream PTBP1 binding site. On the other hand, PTBP2, which is strongly induced in the *Ptbp1* KO cell line and is seen to bind pri-miR-124-1 *in vitro* and in iCLIP analyses of brain tissue (data not shown), does not inhibit miR-124 expression. PTBP1 and PTBP2 have very similar RNA binding properties yet have different activities in repressing the splicing of certain exons (Markovtsov et al. 2000; Amir-Ahmady et al. 2005; Boutz et al. 2007; Tang et al. 2011; Keppetipola et al. 2012). Thus, it appears that the repression of DROSHA cleavage by PTBP1 involves more than simple binding and occlusion of the cleavage site. It will be interesting to assess how the protein interactions of the two PTB proteins differ when they are bound to pri-miR-124-1.

We identified PTBP1 binding sites in other primary miRNA transcripts, including those of the miR-7-1 allele regulated by HuR and Musashi-2. Unlike the miR-7-1 host transcript, the repression of pri-miR-124-1 processing does not lead to the expression of the host RNA as an mRNA, at least in mESCs. It is possible that in some cells, the pri-miR-124-1 transcript does serve a functional role and that PTBP1 acts to allow its expression. It will be interesting to look at this in the retina, where expression of pri-miR-124-1 RNA is observed earlier than the mature miRNA, possibly due to PTBP1 repression (Sanuki et al. 2011).

## Materials and Methods

### Cell lines and tissue culture

The N2a neuroblastoma cell line was grown in DMEM (Fisher Scientific) supplemented with 10 % FBS (Omega Scientific). The mESC line, E14 (Hooper et al. 1987), *Ptbp1*^loxp/loxp^ WT and KO mESCs, were cultured on 0.1 % gelatin-coated dishes with mitotically inactivated mouse embryonic fibroblasts (MEF) (CF1, Applied StemCell, Inc.) in ESC media. ESC media consisted of DMEM (Fisher Scientific) supplemented with 15 % ESC-qualified fetal bovine serum (Thermo Fisher Scientific), 1x non-essential amino acids (Thermo Fisher Scientific), 1× GlutaMAX (Thermo Fisher Scientific), 1× ESC-qualified nucleosides (EMD Millipore), 0.1 mM β-Mercaptoethanol (Sigma-Aldrich), and 10^3^ units/ml ESGRO leukemia inhibitor factor (LIF) (EMD Millipore). Mouse primary cortical cultures were prepared from gestational day 15 C57BL/6 embryos (Charles River Laboratories), as described previously (Zheng et al. 2010). Briefly, cortices were dissected out into ice cold HBSS and dissociated after a 10 min digestion in Trypsin (Invitrogen). Cells were plated at 5 million cells per 78.5 cm^2^ on 0.1 mg/ml poly-L-lysine coated dishes and cultured for 5 days. Neurons represented 70 – 80 % of the cells in the culture. A mouse neuronal progenitor cell line was established from cortical cells of gestational day 15 embryos generated by crossing homozygous Nestin-GFP transgenic mice to wild type C57BL/6. GFP positive cells were collected using FACS and plated on uncoated culture dishes. These NPCs were grown in DMEM/F12 supplemented with B27 (without vitamin A, Thermo Fisher Scientific), 1 mM GlutaMax and antibiotics. EGF and FGF (PeproTech) were added every day at 10 ng/ml concentration. All experiments were approved by the UCLA Institutional Animal Care and Use Committee (ARC& 1998-155-53).

### Transfection and protein/RNA extraction

The plasmids pAAV-nEF-tRFP and pAAV-nEF-tRFP-p2A-FLAG-PTBP1.4 were transfected into mESC (*Ptbp1* WT and KO cell lines) using Lipofectamine 2000 (Thermo Fisher Scientific). 48 hrs post-transfection, protein was extracted with RIPA buffer (50 mM Tris-Cl [pH8.0], 150 mM NaCl, 1 % Igepal CA-C630, 0.5 % Sodium dexolycholate, 0.1 % SDS, 1× Phosphatase Inhibitor, 1× Protease Inhibitor) and RNA was extracted using Trizol (Thermo Fisher Scientific).

### Immunoprecipitation

mESC (E14) were harvested and sonicated in cold Buffer D-200K (20 mM Tris-Cl [pH8.0], 200 mM KCl, 0.2 mM EDTA). After centrifugation at 14,000 rpm for 15 min at 4^°^C, the supernatant was incubated with anti-PTBP1 (PTB_NT) or rabbit IgG isotype control (Thermo Fisher Scientific) at 4^°^C. After 1 hr, proteinG sepharose beads (GE Healthcare) were added to the reaction and further incubated for 1hr. The beads were washed four times in BufferD-200K, and RNA was extracted with phenol.

### RNA isolation, RT-PCR, RT-qPCR and quantification

Total RNA was collected from cell cultures using Trizol (Thermo Fisher Scientific). RNA was treated DNase I (Takara) followed by phenol extraction. 0.8 - 1 µg of total RNA was used for random priming in a 10 µl reaction with 100 units of SuperScript III RT (Thermo Fisher Scientific). PCR was performed using Phusion DNA polymerase (Fisher Scientific). PCR conditions are described in the Supplemental Materials and Methods. The miR-294 and miR-295 stem-loops are located 48 nt apart within the same precursor RNA. This precursor is denoted as miR-294/295. RT-PCR products were run on agarose gels with Ethidium Bromide staining, visualized under UV light, and the band intensities were measured using ImageJ. RT-qPCR was performed using the SensiFAST SYBR Lo-ROX Kit (Bioline) on a QuantStudio 6 Real-Time PCR System (Thermo Fisher Scientific). 1 µg of total RNA was used to generate cDNA for quantification with gene specific primer. Absolute RNA levels were determined using standard curves generated with known amounts of synthesized cDNAs (Bustin 2000). Mature-miRNA reverse transcription and qPCR were performed using TaqMan MicroRNA Reverse Transcription kit and TaqMan Universal PCR Master Mix (Thermo Fisher Scientific, Supplemental Table S3). Synthesized small RNA was used to make a standard curve for absolute quantification. A list of primer sequences is presented in Supplemental Table S2.

### *In vitro* RNA pull-down assays

The RNA pull-down assay was modified from Heo et al. (Heo et al. 2009). WT and chimeric pri-miR-124-1s were prepared by *in vitro* transcription from PCR product templates (Supplemental Materials and Methods). 5’-biotinylated adapter DNAs complementary to the 3’ extensions of the transcripts were used to immobilize the WT and chimeric pri-miR-124-1. Adapter DNAs were first incubated with streptavidin-conjugated sepharose beads (GE Healthcare) in Buffer I-200K (20 mM Tris-Cl [pH 8.0], 200 mM KCl, 2 mM EDTA [pH8.0]) for 1 hr at 4^°^C. Empty streptavidin binding sites were then blocked with free biotin for 10 min. The beads were washed twice with Buffer D-300K (20 mM Tris-Cl [pH 8.0], 300 mM KCl, 0.2 mM EDTA) and then incubated with pri-miRNA in Buffer D-200K with 100 units/ml of RNase inhibitor (Thermo Fisher Scientific) for 2 hrs at room temperature. The beads were washed twice with Buffer D-300K and once with Buffer D-200K. mESC were collected and sonicated in Buffer D-200K. After clearing cell debris by centrifugation, total cell extract was added to the immobilized pri-miRNAs and incubated with constant rotation for 2 – 3 hrs at 4^°^C. After washing with Buffer D-200K twice, proteins were isolated in SDS buffer (50 mM Tris-Cl [pH 6.8], 0.05 % Bromophenol blue, 10 % Glycerol, 2 % SDS and 0.1 M DTT), and separated by 8 % SDS-PAGE for western blotting. Western blot images were taken on a Typhoon imager. RNA was extracted from the beads in phenol, run on standard urea gels, stained with SYBR gold (Thermo Fisher Scientific), and imaged on a Typhoon Imager (GE Healthcare). Fluorescent band intensities of western blots and RNA gels were quantified with ImageQuant software (GE Healthcare).

### PTBP protein quantification

Whole-cell lysates (WCL) were extracted in RIPA, and quantified with the bicinchoninic acid (BCA) method (Smith et al. 1985). 12 µg of WCL protein was loaded on 10 % SDS-PAGE adjacent to a standard curve of His-PTBP recombinant protein. PTBP bands were detected with anti-PTBP1 (PTB_NT) and anti-PTBP2 (PTB_IS2) (Supplemental Table S4), and fluorescent (Cy3) secondary antibodies and imaged on a Typhoon Imager. Band intensities were quantified with ImageQuant software (GE Healthcare).

### Generation of WT and *Ptbp1* KO mESC

A *Ptbp1* mouse line (*Ptbp1*^loxp/loxp^) carrying loxp sites flanking *Ptbp1* exon 2 was obtained from the Xiang-Dong Fu lab (UCSD, generation of this mouse will be described elsewhere) and crossed to an actin-flip recombinase mouse line to remove the neomycin selection cassette in intron 2. *Ptbp1*^loxp/loxp^ mESCs were generated from this *Ptbp1*^loxp/loxp^ mouse following a previously published protocol (Czechanski et al. 2014). Briefly, healthy late blastocysts were collected at 3.5 days post conception (dpc) and cultured overnight in KSOM medium (DMEM high glucose with 15 % KO serum, 2 mM GlutaMAX, 1 mM sodium pyruvate, 0.1 mM MEM NEAA, 0.1 mM β-Mercaptoethanol, 10^3^ units/ml of LIF, 1 µM PD0325901 and 3 µM CHIR99021). Blastocysts were transferred to defined serum-free medium (1 : 1 mixture of DMEM-F12/N2 and Neurobasal/B27, 1 mM GlutaMAX, 0.5 mM sodium pyruvate, 0.1 mM MEM NEAA, 0.1 mM β-Mercaptoethanol) plus 2i/LIF (EMD Millipore) to allow attachment to the feeder layer. After outgrowth from the blastocysts, cells were disaggregated and cultured in the serum-free medium plus 2i/LIF together with MEF. Cells were monitored daily for colony formation. Cells exhibiting differentiation or atypical ES cell morphology were discarded and the healthy ES cell lines were maintained in the serum-free medium plus 2i/LIF with MEF. These *Ptbp1* ^loxp/loxp^ WT mESCs were used to generate *Ptbp1* KO mESCs by transduction with Cre-GFP lentivirus for 3 days. FACS selected GFP positive cells were cultured and genotyped to confirm homozygous deletion of *Ptbp1* exon 2.

### Subcellular fractionation, RNA sequencing and data analysis

These datasets will be described in detail elsewhere. Briefly, total Ribo-minus RNA was isolated from mESCs (clone E14) that were fractionated into cytoplasmic, soluble nuclear, and chromatin pellet compartments as described previously (Wuarin and Schibler 1994; Pandya-Jones and Black 2009; Yeom and Damianov 2017). Paired-end libraries were constructed using the TruSeq Stranded mRNA Library Prep Kit (Illumina). The libraries were subjected to 75 nt paired-end sequencing at the UCLA Broad Stem Cell Center core facility on an Illumina HiSeq2000 to generate 200–250 million mapped reads per sample. Sequences were aligned using STAR (STAR_2.5.1b) to examine the miR-124-1 locus. Further analysis of these datasets is ongoing. The iCLIP data are described in Linares et al. (Linares et al. 2015), and are available at GEO (GSE71179).

Adeno-associated viruses (AAVs) and transduction of cultured cortical neurons AAVs were packaged with serotype 9. AAV2/9 nEF-tRFP and nEF-tRFP-p2A-FLAG-PTBP1.4 were produced by co-transfection of the AAV2 genomic plasmid, pHelper and an AAV9 envelope plasmid into 293FT cells and purified on ioxodianol gradients (Optiprep, Sigma-Aldrich). Virus titer was determined by qPCR. Cultured mouse cortical neurons were transduced with AAV2/9 virus one day after plating (DIV 1), with RNA and Protein then isolated at DIV 8.

### Computational Modeling

All computational simulations were performed in COPASI (Hoops et al. 2006). Ordinary differential equation (ODE) based models were constructed as depicted in Figure 5A as described previously (Mitchell and Mendes 2013). The full system of differential equations with PTBP1 meditated inhibition of miR-124 processing is as follows:

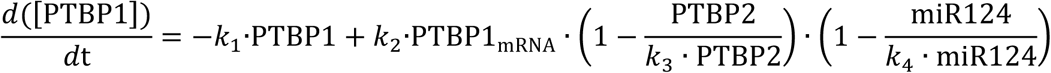

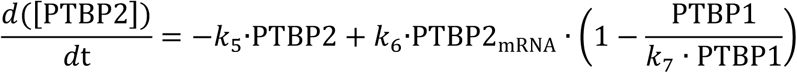

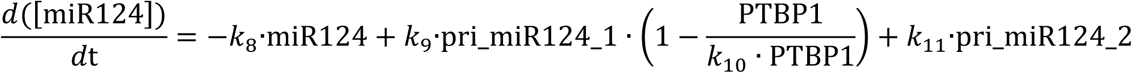

MiR124 denotes the mature miRNA, and pri_miR124_1 and pri_miR124_2 denote the two primary miRNAs. Simulations without PTBP1-meditated inhibition of miR-124 processing were creating by removing the PTBP1 dependent term in Eqn 3 such that pri-miR-124-1 and pri-miR-124-2 processing were modeled in the same way. miRNA and precursor concentrations were experimentally determined as shown in Table 1 and numerically defined. All other parameters were estimated using COPASI parameter estimation to minimize the distance to the remaining molecule numbers in Table 1. Each parameter was constrained between 1 ∙ 10^−6^; and 1 ∙ 10^6^; and particle swarm optimization was performed with a swarm size of 500. Fitting was terminated when the quality of fit no longer improved with repeated iterations (Figure 5B). Repeated single-cell simulations were performed using the “parameter scan” task within COPASI. All kinetic parameters were sampled over a 4-fold range centered on the optimum parameter and a time-course was performed from ESC to the cortical neuron state (Supplemental Figure S5). This process was repeated 1000 times to generate 1000 single-cell simulations with distinct parameters. The range of PTBP2 concentrations within the final population was determined, and cells where PTBP2 was present but below the PTBP1 concentration were measured as “Remaining Cells with [PTBP1]>[PTBP2]” (Figures 5E and F).

## Acknowledgements

We thank Feng Guo (UCLA) for the DGCR8 antibody, Narry V. Kim (Seoul National University, Seoul, South Korea) for the DROSHA-FLAG-pCK plasmid, Grigori Enikolopov (CSHL) for the Nestin-GFP mouse line, and Areum Han, Celine Vuong and Manuel Ares Jr. (UCSC) for critical reading of the manuscript and helpful discussion. This work was supported by National Institutes of Health grant R01 GM049662 to DLB. K-HY was supported by a Postdoctoral Fellowship from Eli and Edythe Broad Center of Regenerative Medicine and Stem Cell Research at UCLA. AJL was supported by the UCLA MSTP program, and the training programs in Neural Repair (T32 NS07449) and Neurobehavioral Genetics (T32 NS048004) at UCLA. SZ was supported by NIH grant R00MH096807. We also acknowledge support from a Quantitative and Computational Biosciences (QCB) Collaboratory Post-doctoral Fellowship awarded to SM and the QCB Collaborator community, directed by Matteo Pellegrini.

## Author contributions

K-HY: Conception and design, Acquisition of data, Analysis and interpretation of data, Drafting and revising the article; SM: Conception and design, Analysis and interpretation of data, Drafting and revising the article; AJL: Contributed essential data; SZ: Contributed essential data and reagents; C-HL: Analysis of data; X-JW: Contributed essential materials; AH: Conception and design, Drafting and revising the article; DLB: Conception and design, Analysis and interpretation of data, Drafting and revising the article.

## Supplemental Figure

**Supplemental Figure S1.**
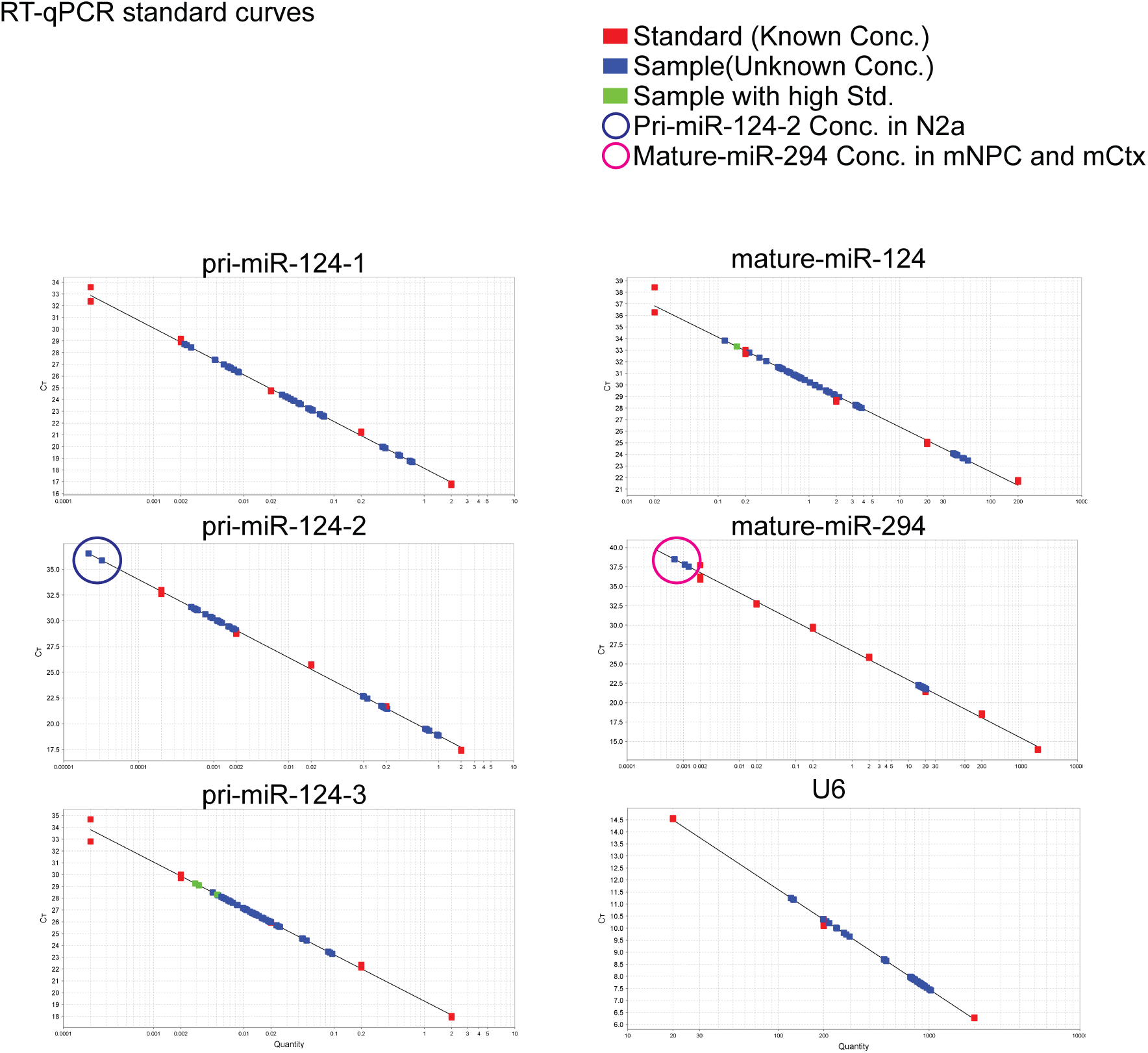
Standard curves used for RT-qPCR quantification. Sample concentrations were determined by comparing Ct values (Y axis) to known concentration standards (X axis). Red dots denote the standard points. Blue dots denote sample points. Pri-miR-124-2 levels in N2a cells were below the lowest standard point and marked with a blue circle. Mature-miR-294 levels in mNPC and mCtx were below the lowest standard point and marked with a magenta circle.

**Supplemental Figure S2.**
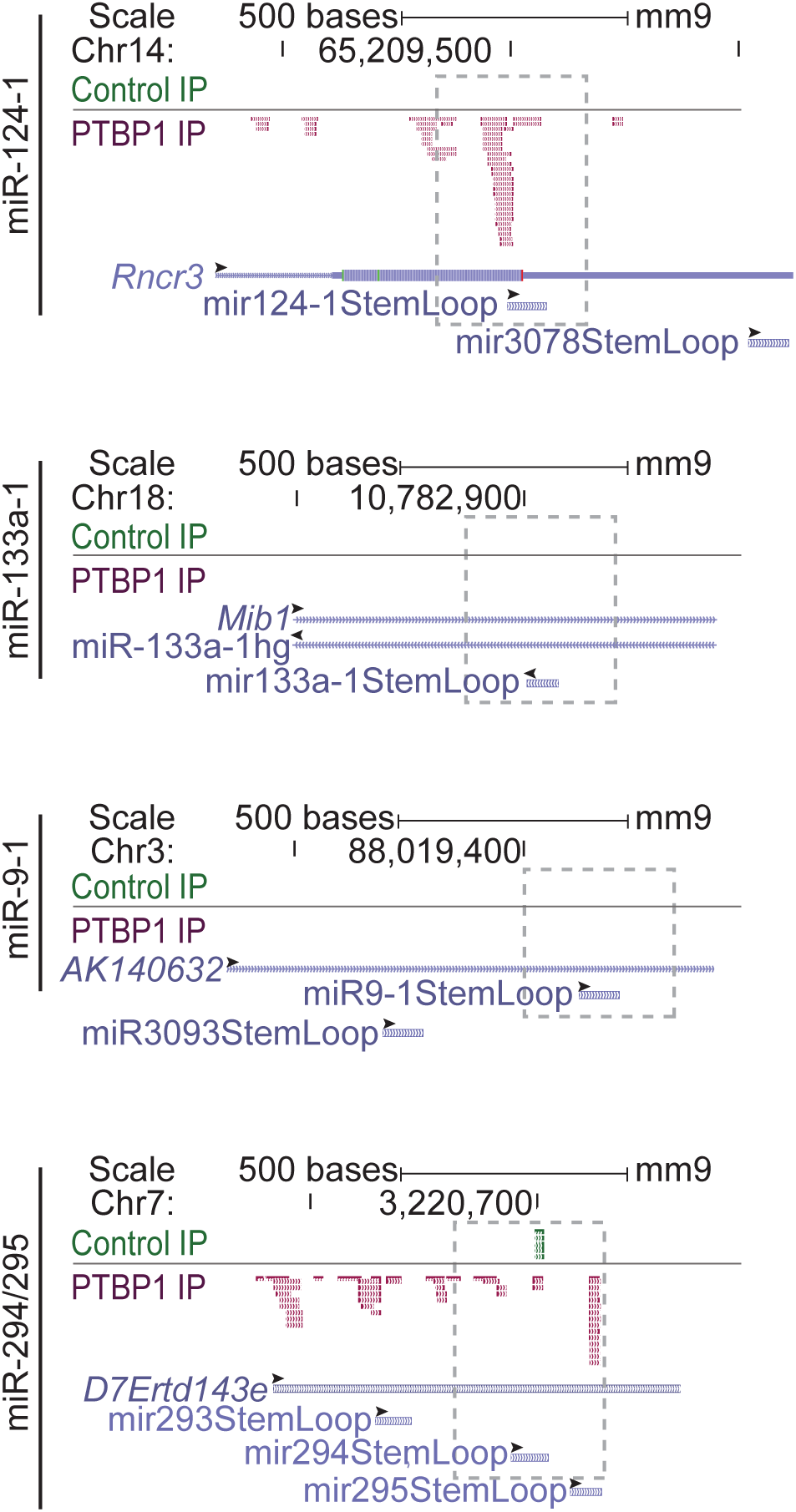
PTBP1 and control iCLIP tags mapping to miRNA loci in mESCs. Genome browser tracks (~ 1 kb window) of loci used in the miRNA PTBP1 IP experiment in Figure 2C. Host transcripts for each miRNA are indicated (light purple). iCLIP tags from control IPs are marked in green and PTBP1 IP iCLIP tags are marked in magenta. Sequence intervals from - 125 nt upstream to + 125 nt downstream of each annotated miRNA stem-loop is marked with a grey dashed box. Arrowheads indicate 5’ to 3’ direction of each of RNA.

**Supplemental Figure S3.**
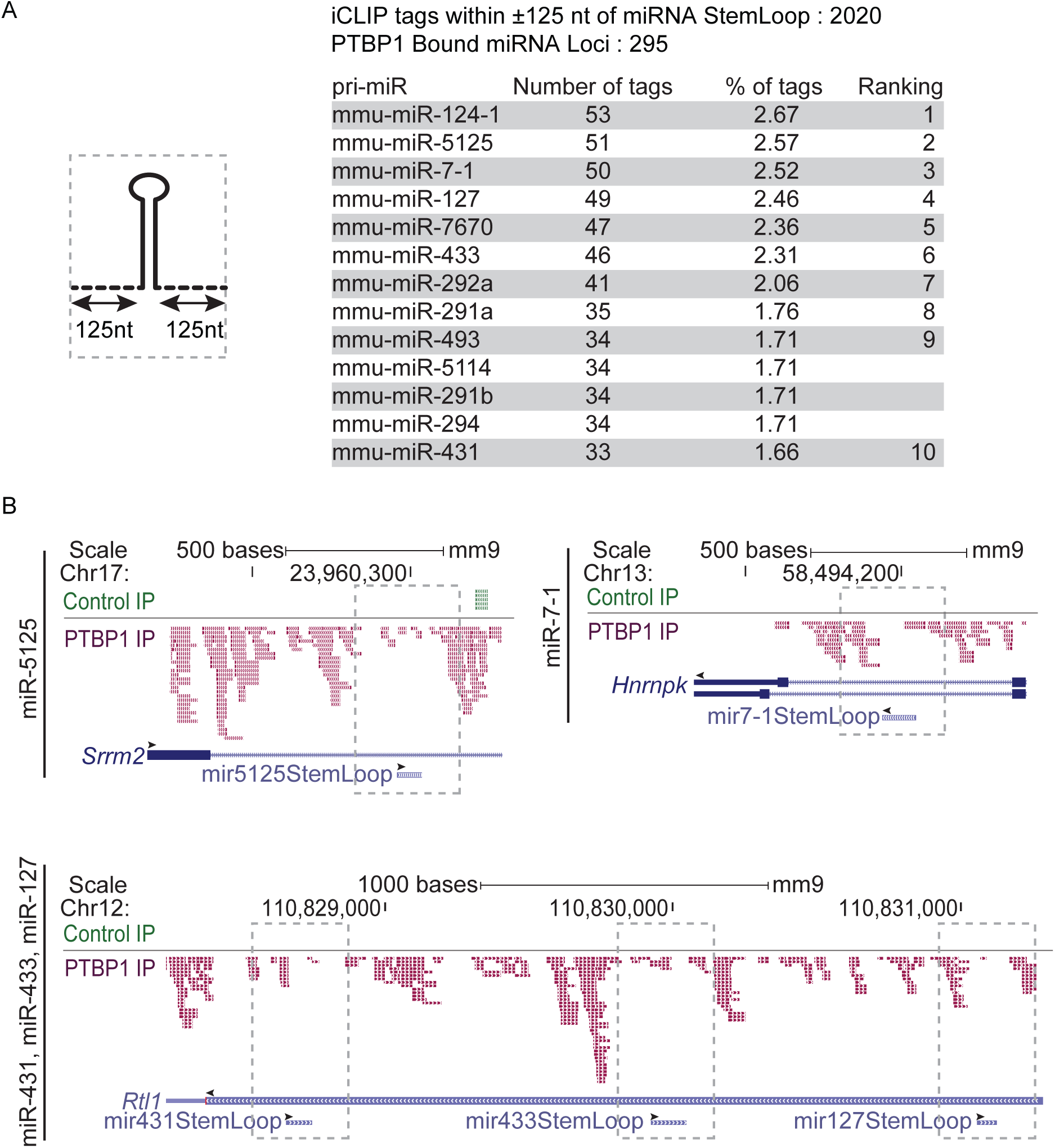
PTBP1 iCLIP tags mapping to miRNA loci in mESCs. A. List of top miRNAs bound by PTBP1. The sequence intervals extending from - 125 nt upstream to + 125 nt downstream of the pre-miRNA stem-loop for all annotated miRNAs (miRbase, Release 21) were assessed for PTBP1 iCLIP tags (left, miRNA stem-loop in solid line; upstream - 125 nt and downstream + 125 nt in dashed line; assessed range for PTBP1 iCLIP tags in the grey dashed box). A total of 2020 PTBP1 iCLIP tags were found to map adjacent to 295 different miRNAs. These pri-miRNAs were ranked by the number of mapped tags and the top 10 are listed on the right panel. B. Genome browser tracks (~ 1 kb and ~ 2 kb window) of miRNA host genes listed in A. Host transcripts for each miRNA are indicated (light purple). iCLIP tags from control IPs are marked in green and PTBP1 IP iCLIP tags are marked in magenta. Assessed range for PTBP1 iCLIP tags are marked by grey dashed box. Several miRNAs show extensive PTBP1 binding at more distal positions than those measured above. Arrowheads indicate 5’ to 3’ direction of each of RNA. Note that for the miR-431 locus at the bottom, miR-431, miR-433 and miR-127 are on the antisense strand, while Rtl1 is a sense transcript. All the PTBP1 iCLIP tags from this locus map in the antisense direction onto the presumptive but not annotated pri-miRNA.

**Supplemental Figure S4.**
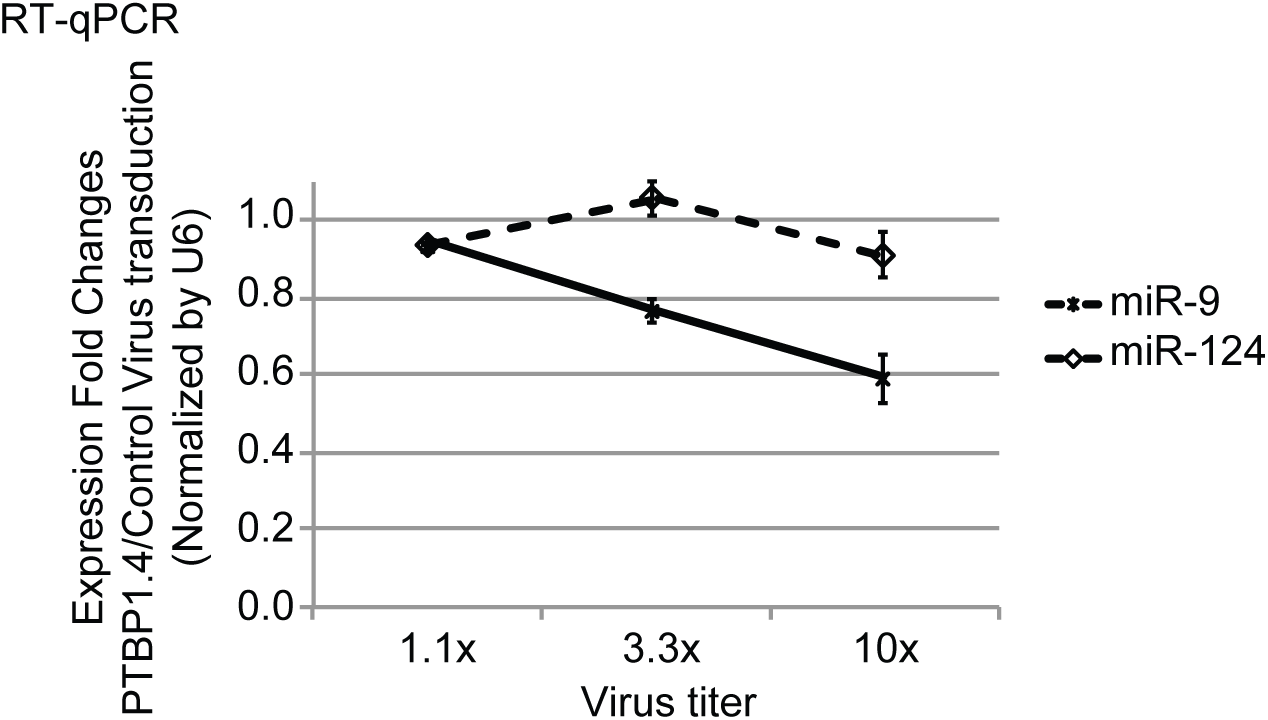
Dependency of miRNA levels on PTBP1-AAV titer in cortical neurons (mCtx) Mature-miRNA levels were measured 8 days after AAV transduction (AAV2/9-nEF-tRFP-p2A-FLAG-PTBP1.4) into cultured cortical neurons. RT-qPCR values were normalized to U6 and to the control transduction (AAV2/9-nEF-tRFP). (n=2 biological replicates; error bars are s.e.m.)

**Supplemental Figure S5.**
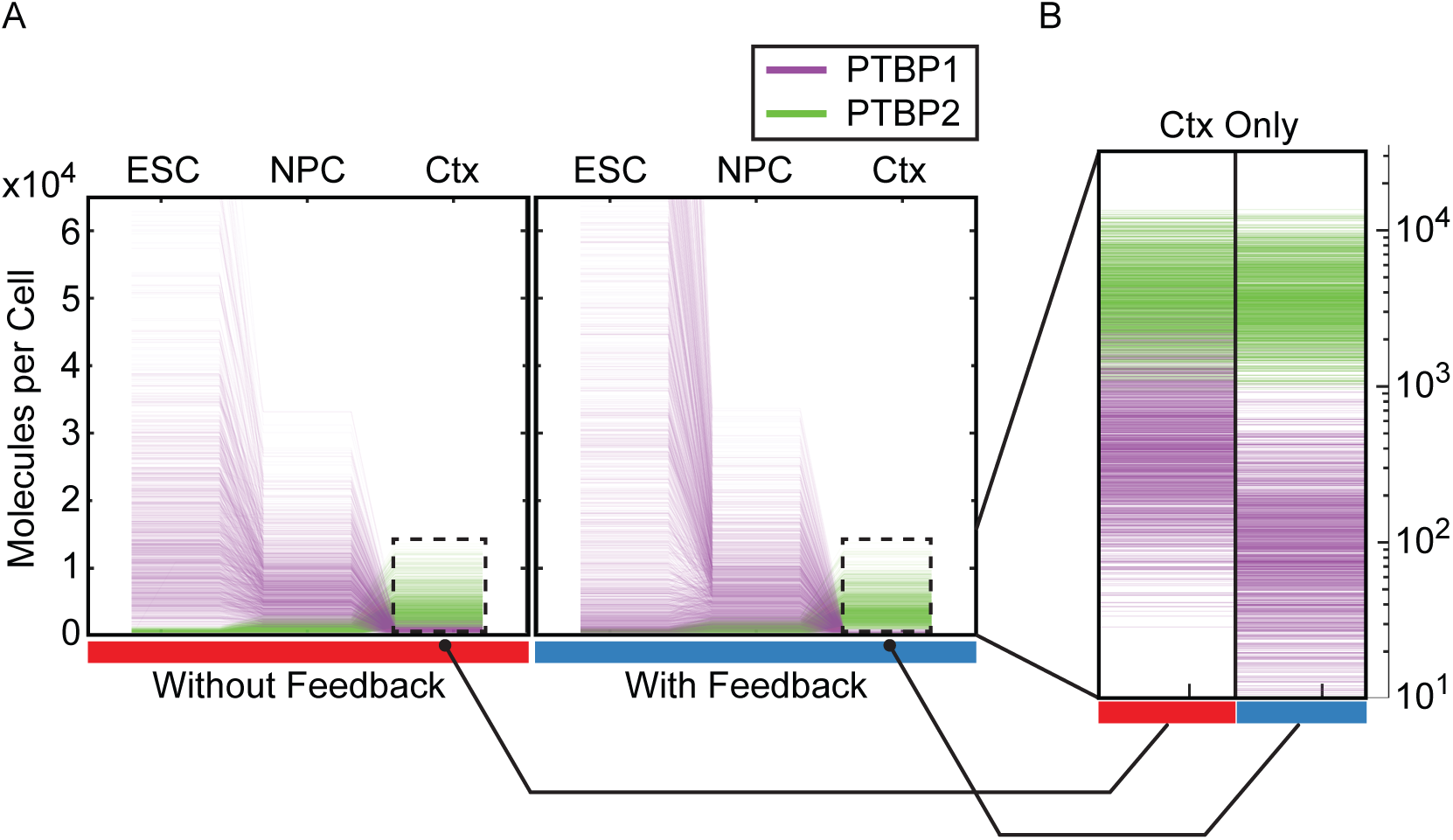
Dependency of miRNA levels on PTBP1-AAV titer in cortical neurons (mCtx).Single-cell model time-courses with distributed parameters. A. PTBP1 and PTBP2 concentrations during modeled single cell differentiation from embryonic stem cell (ESC) to neuronal progenitor cell (NPC) to cortical neuron (Ctx). PTBP1 (purple) and PTBP2 (green) concentrations in 1000 single-cell simulations in the model without inhibition of miR-124-1 processing (left, red bar) and with inhibition of miR-124-1 processing (right, blue bar) during a time-course simulation of neuronal differentiation from ESC through to NPC and Ctx. Parameters were sampled from a distribution centered on the optimal parameter set identified by parameter estimation (Figures 5A and 5B) with maximum and minimum covering a 4-fold range around this parameter. B. Expanded view of PTBP1 and PTBP2 concentrations in Ctx in the model without and with inhibition of miR-124-1 processing.

**Supplemental Table S1.** Quantification of total RNA and total protein per cell. Total RNA was extracted from cells, treated with DNaseI and measured by Nanodrop spectrophoto-meter. Total protein was extracted in RIPA buffer followed by protein BCA quantification. RNA and protein values were divided by cell number, as determined by hemacytometer counting. N2a, mouse neuroblastoma cells; mESC, mouse embryonic stem cells; mNPC, mouse neuronal progenitor cells; mCtx, mouse cortical neurons. The mean ± s.d. of 3 cultures are given.

|  | N2a | mESC | mNPC | mCtx |
| --- | --- | --- | --- | --- |
| Total RNA (pg) per cell | 11.9 ± 0.5 | 19.1 ± 0.8 | 10.8 ± 0.5 | 10.4 ± 0.6 |
| Total Protein (ng) per cell | 0.52 ± 0.03 | 0.37 ± 0.02 | 0.44 ± 0.01 | 0.46 ± 0.03 |

**Supplemental Table S2.**
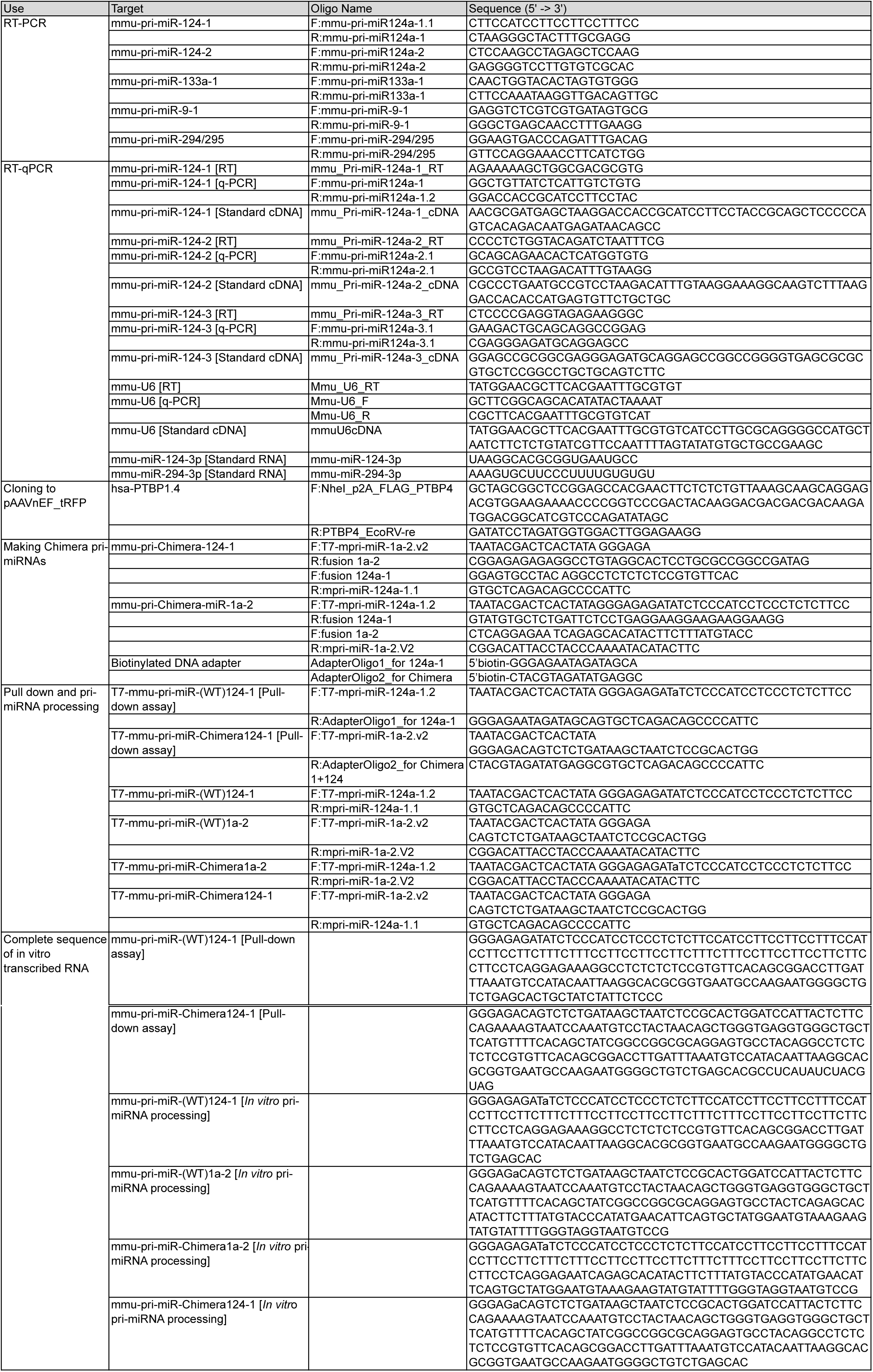
Sequences of oligos and primers used in this study.

**Supplemental Table S3.** List of TaqMan miRNA assays used in this study.

Yeom\_Supplemental Table S3 List of TaqMan miRNA assays used in this study.
| Use | Target | miRBase ID/ NCBI Accession ID | Assay ID (Thermo Fisher) |
| --- | --- | --- | --- |
| mature-miRNA-RT-qPCR | mmu-miR-124a | mmu-miR-124-3p | 4427975-001182 |
|  | mmu-miR-294 | mmu-miR-294-3p | 4427975-001056 |
|  | mmu-miR-9 | mmu-miR-9-5p | 4427975-000583 |
|  | mmu-let7-e | mmu-let-7e-5p | 4427975-002406 |
|  | U6 snRNA | NR_004394 | 4427975-001973 |

**Supplemental Table S4.** List of antibodies used in this study.

Yeom\_Supplemental Table S4 List of antibodies used in this study.
| Target | Antibody name (host/Source ) | Used in |
| --- | --- | --- |
| PTBP1 | PTB_NT (Rabbit/DB lab made) | W/B, Absolute Quantification and PTB-IP |
| PTBP2 | PTB_IS2 (Rabbit/DB lab made) | W/B, Absolute Quantification |
| PTBP1/2 | PTB_CT (Rabbit/DB lab made) | W/B |
| DGCR8 | NC1 (Rabbit/FG lab made) | W/B |
| GAPDH | GPADH 6C5 (Mouse/Thermo Fisher Scientific, AM4300) | W/B |
| U1-70K | SNRNP70 (Rabbit/DB lab made) | W/B |
| alpha-tubulin | alpha-tubulin DM1A(Mouse/Millipore, CP06) | W/B |
| Histone H3 | Histone H3 (Rabbit/Abcam, ab1791 ) | W/B |
| - | Rabbit IgG isotype Control (Rabbit/Thermo Fisher Scientific, 02-6102) | W/B and Control-IP |
| rabbit IgG light chain | HRP conjugated Anti-Rabbit IgG, Light Chain Specific (Mouse/Jackson ImmunoResearch, 211-032-171) | W/B (Figure 2C) |

